# Informative neural representations of unseen contents during higher-order processing in human brains and deep artificial networks

**DOI:** 10.1101/2021.01.12.426428

**Authors:** Ning Mei, Roberto Santana, David Soto

## Abstract

A framework to pinpoint the scope of unconscious processing is critical to improve our models of visual consciousness. Previous research observed brain signatures of unconscious processing in visual cortex but these were not reliably identified. Further, whether unconscious content is represented in high-level stages of the ventral visual stream and linked parieto-frontal areas remains unknown. Using a within-subject, high-precision fMRI approach, we show that unconscious contents can be decoded from multivoxel patterns that are highly distributed alongside the ventral visual pathway and also involving parieto-frontal substrates. Classifiers trained with multivoxel patterns of conscious items generalised to predict the unconscious counterparts, indicating that their neural representations overlap. These findings suggest revisions to models of consciousness such as the neuronal global workspace. We then provide a computational simulation of visual processing/representation without perceptual sensitivity by using deep neural networks performing a similar visual task. The work provides a framework for pinpointing the representation of unconscious knowledge across different task domains.

## Introduction

The neuroscience of consciousness aims to explain the neurobiological basis of subjective experience - the personal stream of perceptions, thoughts and beliefs that make our inner world. This requires a sound framework for addressing the scope of unconscious information processing. Influential neurocognitive models of visual consciousness such as the global neuronal workspace model propose that conscious awareness is associated with sustained activity in large-scale association networks involving fronto-parietal cortex, making information globally accessible to systems involved in working memory, report and behavioural control^1^. Unconscious visual processing, on the other hand, is thought to be transient and operate locally in domain-specific systems -i.e. supporting low-level perceptual analysis^2^. Recent studies have however confronted this view with intriguing data suggesting that unconscious information processing is implicated in higher-order operations associated with cognitive control^3^, memory-guided behaviour across both short- and long-term delays^4–8^, and also language computations^9^; however, follow-up work did not support this view^10^, and even the evidence for unconscious semantic priming has been recently called into question^11,12^. The limits and scope of unconscious information processing remain to be determined.

This controversy is likely to originate from the lack of a sound framework to isolate unconscious information processing^13^. Studies often rely only on subjective measures of (un)awareness^14^ to pinpoint the neural markers of unconscious processing, but these measures are sensitive to criterion biases for deciding to report the presence or absence of awareness^15^, hence making it impossible to determine whether “subjective invisibility” is truly associated with unconscious processing. Previous studies reported brain signatures of unconscious contents in visual cortex^16–19^, but these signatures have not been identified in a reliable manner^20–23^. In these studies using objective measures of (un)awareness, perceptual sensitivity tests are collected off-line, outside the original task context, and typically employ a low number of trials per participant to conclusively exclude conscious awareness and meet the null sensitivity requirement^24,25^. This approach to study unconscious information processing is therefore limited.

Here we used a high-precision, highly-sampled, within-subject approach to pinpoint the neural representation of unconscious contents, even those associated with null perceptual sensitivity, by leveraging the power of machine learning and biologically plausible computational models of visual processing. Critically, the extent to which unconscious content is represented in high-level processing stages along the ventral visual stream and linked prefrontal areas^26^ remains unknown. Previous functional MRI studies indicate the role of conscious awareness in this regard; object categories of visible stimuli are represented in ventral-temporal cortex^27–29^ and parieto-frontal cortex is involved in the representation of conscious perceptual content^30–32^. Here we used a high-precision fMRI paradigm to contrast these views. We further asked the extent to which the representation of unconscious content maps onto the representations of the conscious counterparts. This issue remains unsolved^16,33^, yet it has ramifications for models of consciousness such as the neuronal global workspace^34^.

Subsequently, we used deep feedforward convolutional neural network models (FCNNs)^35–37^ to provide a representational level^38^ simulation of visual representations/processing in the absence of perceptual sensitivity. FCNNs were used given their excellent performance in image classification^35,39,40^ and given the known similarities between the representational spaces during object recognition in FCNNs and high-level brain regions in ventral visual cortex^41,42^. FCNNs performed the same task given to the human participants using the same images corrupted by different levels of noise. We asked whether, similar to the brain, the animate vs inanimate dimension of the stimulus could be decoded by analysing the activity state of the hidden layer of the FCNN network, despite the fact that the network itself had no perceptual sensitivity at identifying the image class.

## Results

Observers (N = 7) performed six fMRI sessions across 6 days leading to a total of 1728 trials per subject, allowing us to pinpoint meaningful and reliable neural patterns of conscious and unconscious content within each observer. Observers were presented with gray-scaled images of animate and inanimate items with a random-phase noise background^43^. The target images were presented briefly, preceded and followed by a dynamic mask composed of several frames of Gaussian noise (Figure 1). On each trial of the fMRI experiment, participants were required to discriminate the image category and to indicate their subjective awareness (i.e. (i) no experience/just guessing (ii) brief glimpse (iii) clear experience with a confident response). The duration of the images was based on an adaptive staircase that was running throughout the experiment (see Methods), and which, based on pilot testing, was devised to obtain a high proportion of unconscious trials. On average, target duration was (i) 25 ms on trials rated as unaware (ii) 38 ms on glimpse trials and (iii) 47 ms on the aware trials. A one-way repeated measures ANOVA showed a significant effect of visibility on the target’s duration (F(2,12) = 321.98, p < 0.001, *η*^2^ = 0.98). Due to the online adaptive staircase that was running throughout the experiment to achieve null sensitivity on the unconscious trials, the signal to noise ratio of the image was higher in the conscious trials. Of note, if stimulus parameters were kept constant throughout the experiment it would have been impossible to obtain trials associated with null perceptual sensitivity alongside the conscious trials. Our paradigm was not designed to assess the neural correlates of consciousness^44^ (i.e. the difference between unseen and seen items), because differences in neural activity between unseen and seen conditions might be partially due to differences in stimulus duration. However, our paradigm addresses the similarities between conscious and unconscious representations. This is done by testing whether a pattern classifier trained in the visible trials generalised to predict the category of unconscious items. Our primary goal, however, was to provide a precise and conservative assessment of the neural representation of unseen contents.

**Figure 1:**
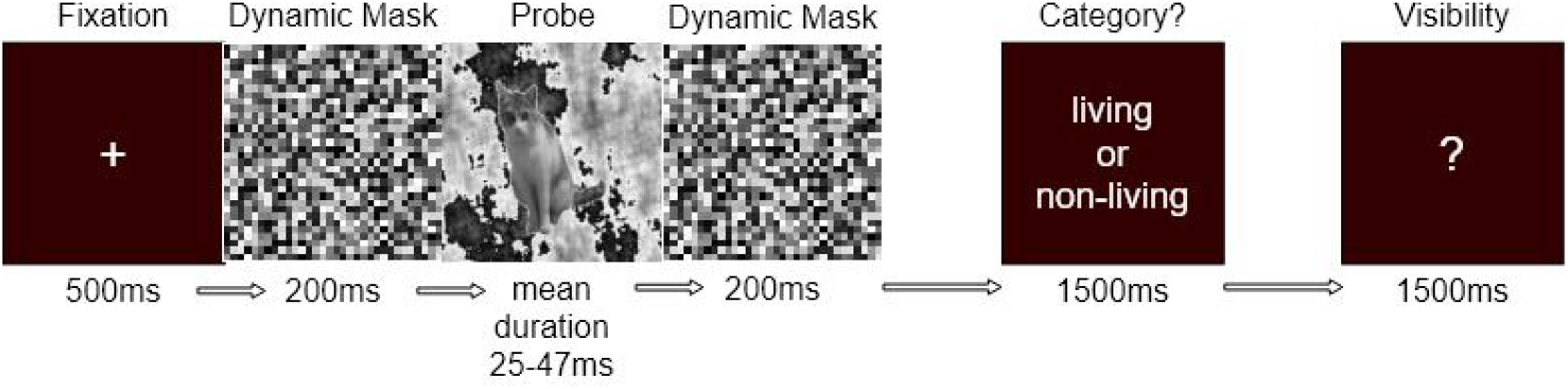
Example of the sequence of events within an experimental trial. Observers were asked to discriminate the category of the masked image (living vs. non-living) and then rate their visual awareness on a trial by trial basis. Example of a trial with a masked image of a cat. The stimuli were selected from an open source image set^43^.

### Behavioral performance

First, we assessed whether observers’ performance at discriminating the image category was above chance in each of the awareness conditions by using a signal detection theoretic measure to index perceptual accuracy, namely, the non-parametric A’^45^. Permutation tests were performed to estimate the empirical chance level within each observer (see Methods). All observers displayed above chance perceptual sensitivity in both glimpse and visible trials (p < 0.001, permuted p-values). Four of the seven subjects showed null perceptual sensitivity (*p*_*sub*–01_ = 0.44, *p*_*sub*–02_ = 0.11, *p*_*sub*–03_ = 0.54, and *p*_*sub*–04_ = 0.20) in those trials in which participants reported a lack of awareness of the images. Discrimination performance in two additional participants deviated from chance (*p*_*sub*–05_ = 0.03, *p*_*sub*–06_ = 0.02, uncorrected for the number tests across the different awareness states) but only one observer clearly showed above chance performance in the unaware trials (*p*_*sub*–07_ < 0.001). Figure 2 illustrates the distribution of A’ values alongside the chance distribution for each participant. Similar results were obtained using the parametric d’ measure of perceptual sensitivity (see Supplementary Figure 1).

**Figure 2:**
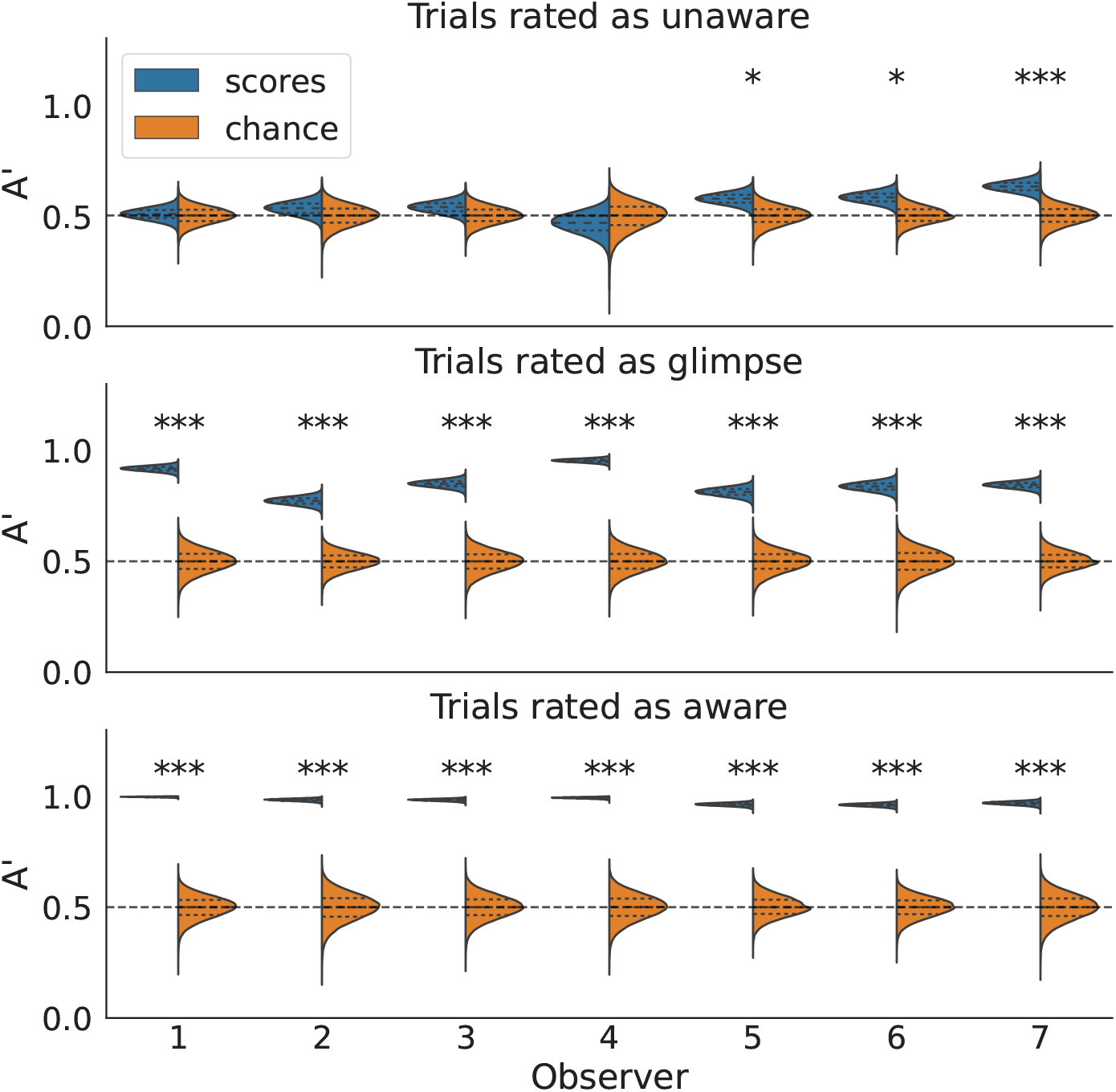
Behavioral performance. Distribution of within-observer A’ scores with mean, first and third quartile, and the corresponding empirical chance distributions for each observer and awareness state. * *p* < 0.05, ** *p* < 0.01, *** *p* < 0.001; one-tailed p-value.

While individual biases in reporting awareness are difficult to control experimentally, several features of the data indicate that participants were using the subjective ratings appropriately. First, the meta-d’ scores that measure how well one’s awareness ratings align with discrimination accuracy were high. Second, as noted above, the average target’s duration set by the adaptive staircase was lowest/highest in the trials rated as unaware/aware. Finally, inspection of the data shows that the proportion of trials rated as aware was similar among the participants showing null sensitivity on the unconscious trials and those showing above chance performance (see Supplementary Table 1).

### FMRI Decoding results

We then used a linear support vector machine (SVM) with out-of-sample generalization to decode the categories of out-of-sample target images in the unconscious and the conscious trials. Trials in which observers reported a ‘glimpse but not confidence in the response’ were not assessed here since we elected to focus on the critical unconscious and conscious trials. The classifier was fed with multivoxel patterns of BOLD responses in a set of 12 a priori regions of interest comprising the ventral visual pathway and higher-order association cortex (see Methods). Permutation tests were run within each subject to estimate the reliability of the decoding at the single subject level (see Methods).

In the unconscious trials, the image class was significantly decoded from activity patterns in visual cortex, including high-level areas in the ventral visual cortex and even in prefrontal regions. Specifically, activity patterns in the fusiform cortex allowed for decoding of unconscious contents reliably within each of the four observers showing null perceptual sensitivity, and moreover the unconscious content could be decoded from prefrontal areas in middle and inferior gyrus in these observers (observers 1 - 4). Figure 3 illustrates the decoding results (see also Supplementary Tables 2, 3 and 4 for summary statistics). A similar pattern was observed in the participants whose perceptual sensitivity deviated from chance (observers 5 - 7). Incidentally, inspection of the distribution of decoding accuracy of the observers that were at chance, and those observers that were above chance in the unconscious trials, did not reveal any advantage for the latter observers despite the unseen contents were used to guide the perceptual decision (see Supplementary Figure 2; but see Stein et al.^46^ for evidence in the visual cortex).

**Figure 3:**
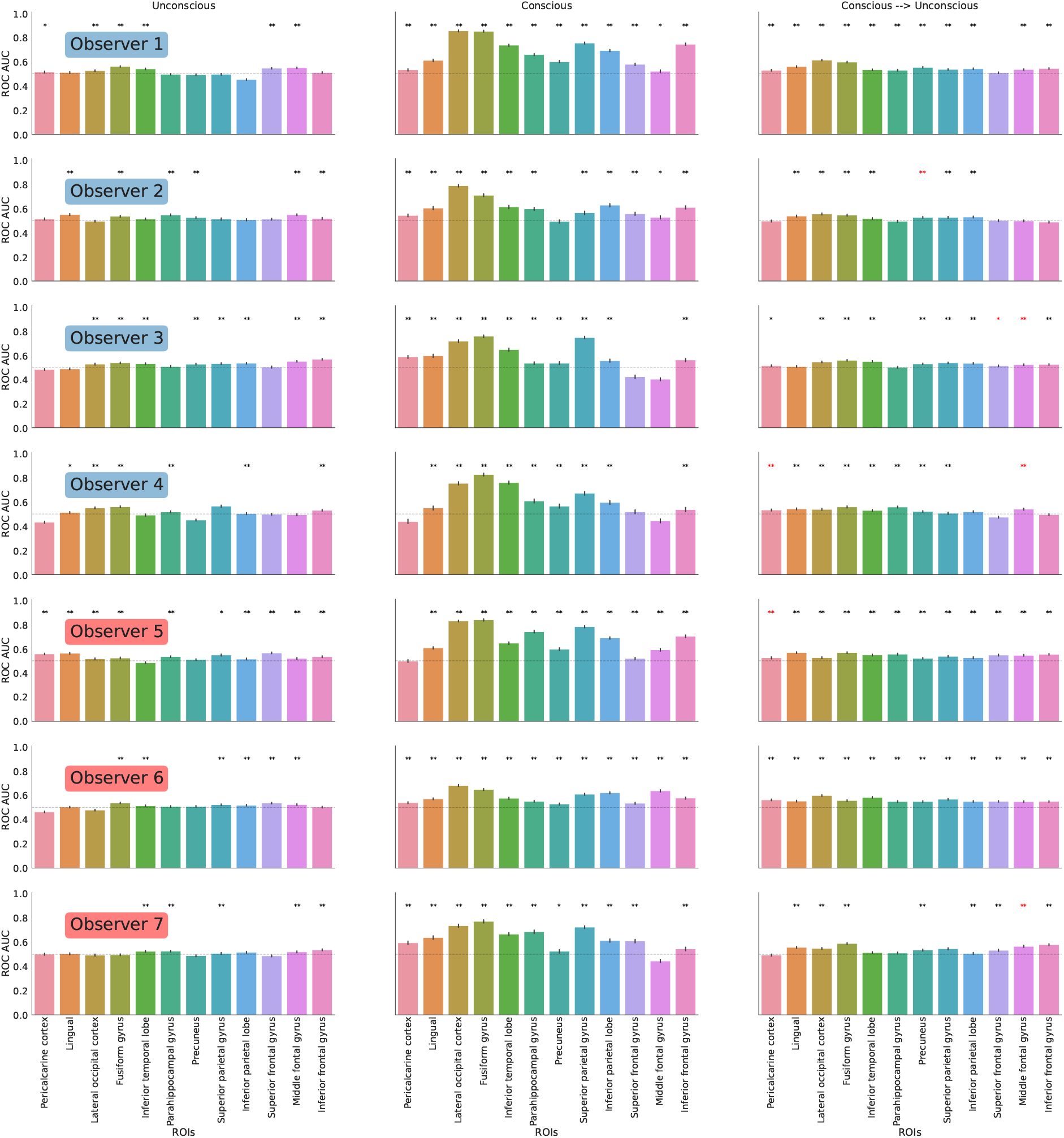
Decoding performance (ROC-AUC) for out-of-sample images for each observer across the unconscious and conscious trials. *:p < 0.05, **:p < 0.01, ***:p < 0.001, one-tailed p-value, after multiple comparison correction for the number of ROIs tested for each observer. Error bars represent the standard error of the mean. Red asterisks indicate those ROIs in which the cross-awareness state generalization appeared above chance but the decoding in the conscious condition was at chance, and these were discarded.

In the conscious trials, the level of decoding accuracy was highly reliable in all subjects tested, across all visual ROIs in the ventral visual pathway, its linked higher-order regions in the inferior frontal cortex^26^, and also inferior and superior parietal cortex. Then, using transfer learning we investigated whether the multivoxel brain representation of perceptual contents in the visible, conscious trials, was similar to those of the unconscious trials. Accordingly, we trained the classifier in the conscious trials and then performed out-of-sample cross-validation in the unconscious trials. The fact that conscious and unconscious stimuli differed in signal strength most likely accounts for the higher classification performance in the conscious trials, but it actually makes the generalization test from conscious to unconscious trials stronger.

The results showed that a decoder trained in the conscious trials using multivoxel patterns in fusiform gyrus, lateral occipital cortex, and precuneus, generalised well to predict the target image in the unconscious trials, remarkably, in all subjects. Also, a decoder trained in the conscious trials with BOLD activity patterns in lingual gyrus, inferior temporal lobe, inferior parietal and superior parietal gyrus generalised to the unconscious trials in 6 out of 7 subjects. We also found that multivoxel patterns in the inferior frontal cortex generalised from conscious to unconscious trials in 5 subjects. Taken together this pattern of results indicates some degree of overlap between the multivariate patterns in conscious and unconscious states across higher-order visual areas and beyond, including parietal and even prefrontal areas in a substantial number of the participants.

We note there were a few cases in which the level of decoding accuracy was below chance. It has been argued that cases of below chance classification may occur when the decoder learns a particular linear mapping between voxels’ activity and the classification targets during the training phase, and the sign of this relationship is reversed in the test set, which may occur with small effect sizes^47^. Computational simulation work^48^, indicated that below-chance classification may be associated with low variance in the distribution of correlations between the voxel activities and the classification targets. In the current study, nevertheless, below-chance classification was not systematic in that critical ROIs showed reliable above chance performance across all subjects. In addition, we verified that the observed pattern of results held across different control analyses. In particular, we repeated the original analysis (i) increasing the L1 regularization term from the default of C = 1 to 5 to try reduce overfitting, and (ii) using a stratified random shuffle split procedure to keep a similar ratio of samples for each class in train and test sets. Supplementary Figures 3 and 4 illustrate these results. Overall the results hold across these follow up analyses, including significant classification accuracy of unconscious contents in higher-order visual areas (i.e. fusiform) and prefrontal areas, while there was little evidence of below-chance classification. However, the generalization performance from conscious to the unconscious trials in prefrontal areas displayed a smaller effect size and was significant in 4-5 participants. It may be possible to improve the generalization from conscious to unconscious trials using domain-adaptation techniques for transfer learning^49^, but this was beyond the scope of the present study.

### Feedforward convolutional neural network model simulations

The key goal of the simulations was to provide a computational framework to study how deep neural networks represent noisy visual items, in particular, those associated with a lack of classification accuracy. Feedforward convolutional neural networks (FCNNs)^35–37,50^ were therefore used to provide a representational model of information processing without perceptual sensitivity. FCNNs were trained on clear images in order to emulate how the brain learns to perform object recognition. FCNNs then performed a similar visual task to the human observers using the same images across different levels of Gaussian noise. FCNNs are known to be excellent in image classification^35,39,40^, but also sensitive to image perturbations^51^; hence, we expected the level of classification accuracy of the FCNNs to drop with increasing levels of noise in the image.

84,000 FCNN model simulations were performed, resulting from combining 5 pre-trained FCNN configurations, 7 hidden layer units, 4 dropout rates, 6 hidden layer activation functions, 2 output layer activation functions, and 50 noise levels (see Methods). Initially, we were interested in those poorly performing FCNNs, and hence we only attempted to decode the stimulus category from the hidden layer in those cases in which the FCNN ROC-AUC classification performance was lower than 0.55. We observed that 61,086 of the FCNN models showed a ROC-AUC score below 0.55 under conditions of increasing noise in the image.

### Informative hidden layer representations in the FCNN models

Then, we asked whether, despite the FCNN failing to classify the image, the animate vs inanimate dimension of the stimulus could still be decoded by analysing the activity patterns of the hidden layer of the network. To test this, a linear SVM was applied to the hidden layer representation for decoding the image class across different levels of noise, even when the FCNN model classification performance was at chance (see Methods). Previous studies modeled visual recognition using FCNN^52,53^, demonstrating that the last hidden layer of FCNNs has representational spaces that are similar to those in high-level regions in ventral visual cortex^52–54^. Therefore, we focused our analyses on the very last hidden layer of the FCNN in the current study, also considering limitations in computational resources due to the large number of simulations (see Methods).

Figure 4^*a*^ shows the classification performance of the FCNN models (black) and also the decoding accuracy SVM applied to the hidden layer representation of the FCNN (blue) as a function of the level of noise and the different factors. When the level of noise was low, FCNN models could classify the category of the images very well reaching ROC-AUC scores higher than 0.9 but performance dropped with the level of Gaussian noise. The observed logarithmic downward trend could be due to the exponential sampling of noise levels (see Methods).

**Figure 4:**
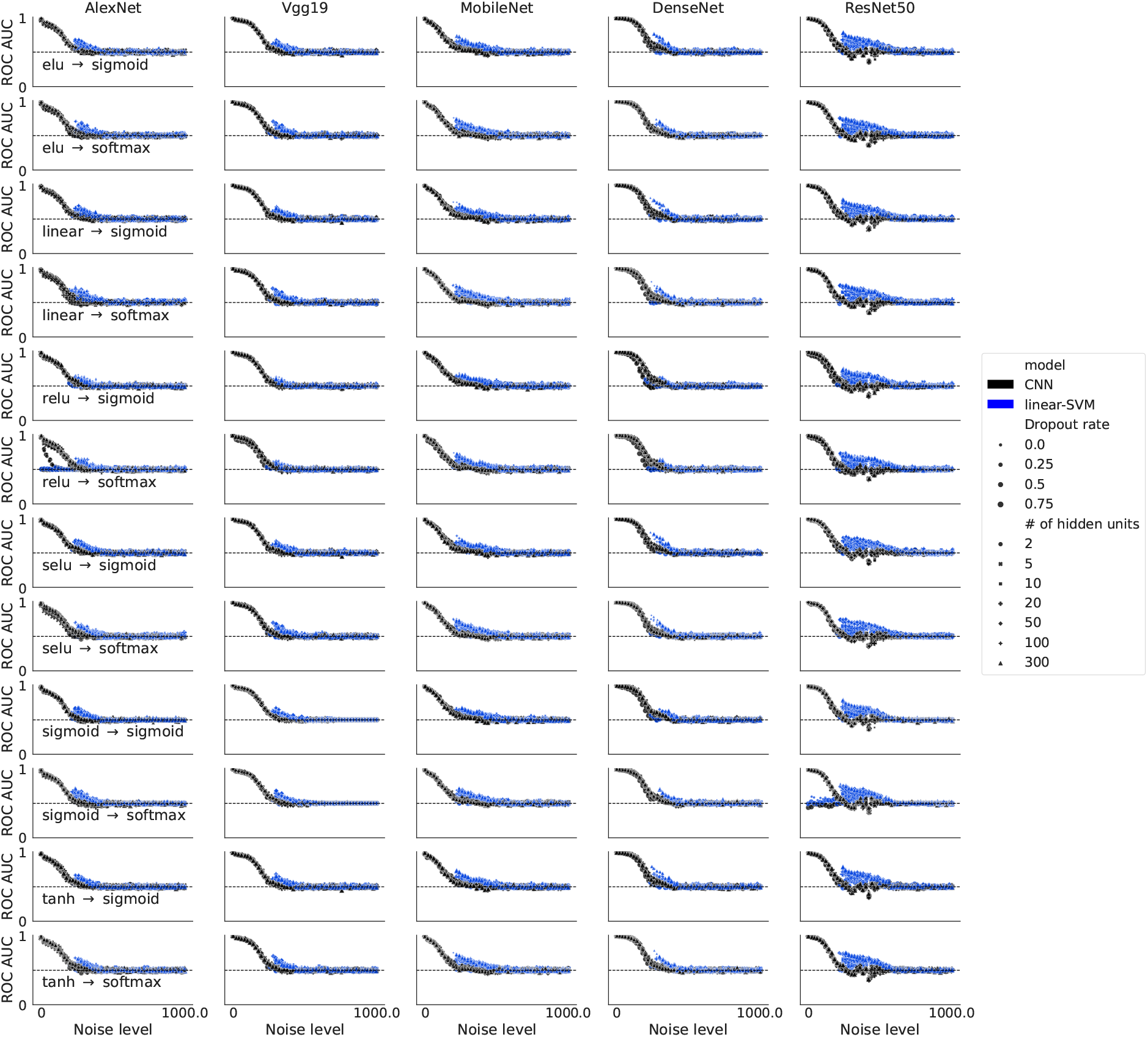
The black dots illustrate the classification performance of the FCNN models, as a function of noise level, type of pre-trained model configuration (column) and activation functions (row). The blue dots illustrate the classification performance of the linear SVMs applied to the hidden layer of the FCNN model when the FCNN classification performance was lower than 0.55.

Remarkably, when the FCNN models failed to classify the noisy images (p > 0.05; N = 32,435), we observed that the hidden layer representation of these FCNN models contained information that allowed a linear SVM classifier to decode the image category above chance levels reliably in 12,777 of the simulations (p < 0.05, one sample permutation test). Figure 5 illustrates the decoding results based on the hidden layer representation when the FCNN was at chance.

**Figure 5:**
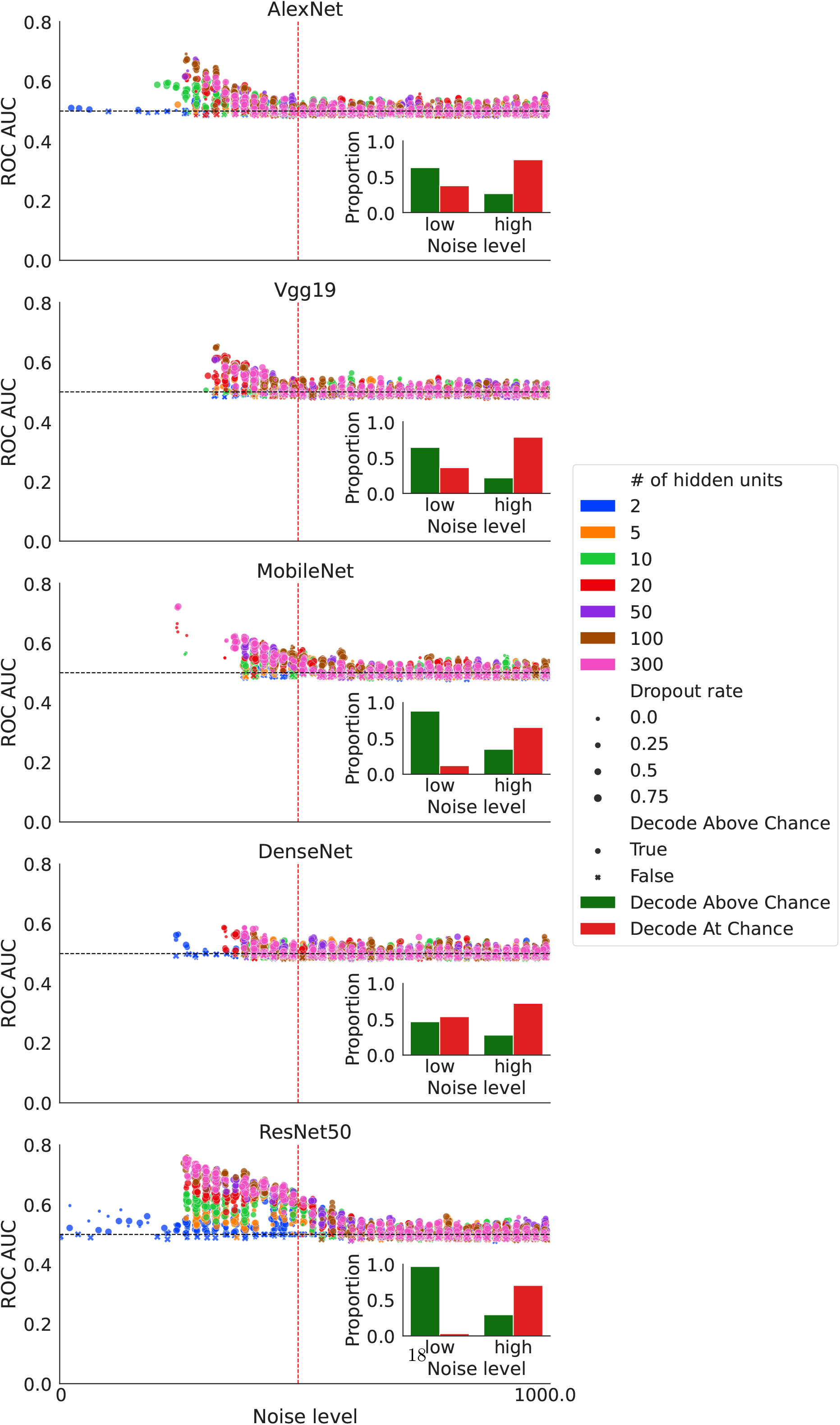

It is noted that even when the noise level was relatively low, some FCNN models such as AlexNet and ResNet did not perform well in the image classification task. Inspection of these models indicated poor performance in the validation phase of the training (prior to testing), which suggests that particular combinations of hidden layer units, activation function, and dropout rate in AlexNet and ResNet impeded learning the classes properly.

When the noise level was relatively low and the FCNN models failed to discriminate the noisy images (N = 7,841), 75.39% of the linear SVMs could decode the FCNN hidden layers and the difference in decoding performance between SVM and FCNN was significantly greater than zero (p < 0.001, one sample permutation test).

Remarkably, even when the noise level was higher and the FCNN classified the images at chance level, 27.91% of the linear SVMs could decode the image category from the FCNN hidden layers. Crucially, the comparison of SVM decoding from the hidden layer and the FCNN classification performance including those 24,584 cases in which the FCNN was at chance again showed a significant difference (permutation p < 0.05), demonstrating that the hidden layer of the FCNN contained informative representations despite the FCNN classification performance was at chance.

MobileNet produced more informative hidden representations that could be decoded by the SVMs compared to other candidate model configurations. We also observed that the classification performance of ResNet models trained with different configurations (e.g., varying number of hidden units, dropout rates) did not fall to chance level until the noise level was relatively high (closer to the dashed line) and the proportion of SVMs being able to decode FCNN hidden layers was higher compared to other model configurations (50.62% v.s. 46.92% for MobileNet, 35.51% for AlexNet, 31.35% for DenseNet, and 30.22% for VGGNet). Additionally, we observed that even when the noise level was high, the MobileNet models provided a higher proportion of hidden representations that were decodable by the linear SVMs (34.77% v.s. 29.53% for ResNet, 27.84% for DenseNet, 26.35% for AlexNet, and 21.62% for VGGNet, see Figure 5). In summary, MobileNet and ResNet50 generated the most informative hidden representations. These networks have an intermediate level of depth. By comparison, the deepest network DenseNet did not produce better hidden representations. Hence the depth of the network per se does not appear to determine the quality or informativeness of the hidden representations significantly.

Then, we sought to further understand the influence of the components of the FCNN architecture (i.e. dropout rate, number of hidden units) on decoding performance. We used a random forest classifier to compute the feature importance of the different FCNN components for predicting whether or not the SVM decoded the image class based on the hidden layer representation. The classification performance was estimated by random shuffle stratified cross-validation with 100 folds (80/20 splitting). In each fold, a random forest classifier was fit to predict whether or not the hidden representation was decodable on the training set, and then the feature importance was estimated by a permutation procedure on the test set^55,56^. Briefly, for a given component (i.e. hidden layer activation function), the order of instances was shuffled while the order of the instances of other components was not changed, in order to create a corrupted version of the data. The dropped classification performance indicated how important a particular feature was. Supplementary Figure 5 shows that the noise level in the image was the best indicator of whether a hidden representation was decodable, followed by model architecture, followed by the number of hidden units, and by the type of hidden activation and output activation functions. The least important feature was the dropout rate. We conducted a one-way ANOVA on the feature importances obtained in each cross-validation fold taking the above features as a factor. This allowed us to quantify the effect of the noise level, model architecture, number of hidden units, type of hidden activation function, type of output activation function, and dropout rate, on the feature importance. There were significant differences between the components of the network models tested (F(5, 594) = 2215.57, p < 0.001, *η*^2^ = 0.95). Post-hoc t tests showed that all the pairwise comparisons were reliable (p < 0.001, two-tailed, Supplementary Table 5, pairwise Tukey-HSD posthoc tests).

We also conducted a control analysis using the SVM to decode the hidden layer representation of clear images. We found that when the images contained little to no noise, the SVM decoding performance was similar to the FCNN performance. However, as the noise level increased, the FCNN performance decreased significantly more than the SVM decoding performance, until they both converged to chance level (see Supplementary Figures 6 and 7). Hence, the significant decoding from the hidden layer was selective to a range of noise levels in the image.

Additional simulations were run to test whether similar results are obtained when the FCNNs are trained with images embedded in Gaussian noise. Here we used a similar pipeline to that used with clear images, except that here the images during training were embedded in Gaussian noise sampled from a standard normal distribution (mean centered at zero with unit variance). The results are shown in Supplementary Figures 8, 9 and 10. To describe the results, here we defined “low noise” as lower than 1 (i.e. the noise level used during training) and “high noise” as greater than 1. In the range of low noise, 98.54% of the VGG19 configurations (91.76% for Resnet50) produced informative hidden representations when the performance of the FCNNs was at chance (p > 0.05 by a permutation-based t test). This was due to the FCNN being sensitive to the removal of the noise when it was tested with clearer images. Recall that FCNNs are known to be very sensitive to image perturbations (Kubilius et al., 2019). Critically, when the level of noise wFas high, 56.40% of the VGG19 configurations (26.67% for Resnet50) produced informative hidden representations when the performance of the FCNNs was at chance (Supplementary Figure 10). The performance of the SVM decoding the hidden layer representation was statistically significant overall across all the noise levels (p < 0.001 in a permutation-based t test, one-tailed), and it was also significant when the noise level was low (p < 0.001, in a permutation-based t test, one-tailed) or high (p < 0.001, in a permutation-based t test, one-tailed).

We then assessed whether the image categories could also be decoded from the first layer activity patterns and we show similar results to those obtained from the hidden layer representations (see Supplementary Materials and Figure 11).

Finally, we attempted to relate the FCNN models to the human brain activity patterns in the unconscious and conscious condition by means of a representational similarity analysis (RSA;^57^). Specifically, RSA measures the similarity of the brain activity patterns across the stimulus space and how they map onto the representation of the images given by the computational model (i.e. the FCNN hidden layer representations of the different stimuli; see Supplementary Information for details on the RSA methods and Supplementary Figures 12 and 13 for a depiction of the model representational dissimilarity matrices). The RSA results indicate that in the unconscious trials, the brain activity patterns were similar to the FCNN models in clusters involving the visual cortex and extending into fronto-parietal areas. In the conscious condition, the similarity between FCNN models and brain activity was higher. The Supplementary Figures 14 and 15 depict the RSA results. Note that these results are however descriptive. The present study was specifically designed to perform high-precision decoding analyses at the single subject level, but we currently lack robust within-subject RSA analytical pipelines to make within-subject statistical inference.

## Discussion

We tested a high-precision, within-subject framework to provide a representational account of the scope of information processing for unseen items, even those associated with null perceptual sensitivity^13^ in both brains and deep artificial neural networks. Isolating the brain representation of unconscious contents has been difficult to achieve in systematic and reliable fashion in previous work^58^, with low numbers of trials and signal detection theoretic constraints^24^ not allowing to decisively discard conscious perception^20,21,23,46,59^, and, critically, when unconscious content could be decoded, this was restricted to visual cortex - see also^22,46^. The current results demonstrate that when human participants and FCNNs models fail to recognise the image content, there remain informative traces of the unseen items in a hidden state of the network during high-level stages of information processing. These hidden representations allow for classification of the animate vs inanimate dimension of unseen perceptual contents. Notably, the fMRI results from our high-precision, highly-sampled, within-subject approach showed that unconscious contents can be reliably decoded from multivoxel patterns that are highly distributed along the ventral visual pathway and also involving prefrontal substrates in middle and inferior gyrus. High-precision fMRI decoding paradigms can thus provide a richer information-based approach^60^ to reveal meaningful feature representations of unconscious content, and that otherwise would be missed. The current findings have implications for models proposing that unconscious information processing is local and restricted to sensory cortex^2^. For instance, according to the neural global workspace model^61^, distributed activity patterns in fronto-parietal cortex are a marker of conscious access^62^. Both the middle frontal gyrus and inferior frontal areas have been implicated in the coding of visible items during working memory tasks^30^ and also in binocular rivalry paradigms used to track moment-to-moment changes in the contents of consciousness^32^. The inferior frontal cortex forms part of the ventral visual pathway that links extrastriate, and inferior temporal areas that is crucial for object recognition^26^. Remarkably, the fMRI decoding results indicate that the neural representation of conscious and unconscious contents overlap in the ventral visual pathway, also including parietal and even to some extent in prefrontal areas in a substantial part of the observers. Previous studies using lowly sampled fMRI designs could not reveal evidence consistent with this view^16,33^. Visual consciousness may be associated with neural representations that are more stable across different presentations of the events^33^, but our data indicates that the underlying representational patterns in terms of perceptual content are to some degree generalizable across awareness states, despite the non-linear dynamic changes in the intensity of the neural response that occur in fronto-parietal cortex during conscious processing,^62–67^, and even despite the visible items were here presented for a longer duration. This observation points to revisions to the neuronal global workspace model^1^.

The generalization of the feature representations across visibility states is also supported by the deep neural network model simulations. FCNNs initially trained with clear visible images, subsequently produced informative feature representations in the hidden layer when they were exposed to noisy images. Prior work showed that FCNNs are a good computational model of the ventral visual pathway^41,42,52,53,68^. FCNNs performed well the perceptual identification task with clear images, also in keeping with prior studies^69–72^. FCNNs are sensitive to image perturbations^51^ and accordingly, FCNNs classification performance dropped as the noise level increased and eventually fell to chance levels. Crucially, in these conditions, the hidden representation of the FCNNs contained informative representations of the target class despite the classification accuracy was at chance.

A critical issue is whether the informative traces of the unseen contents found in both brains and deep artificial networks can be understood in terms of representational account. The notion of neural representation may be based on a functional perspective -as conveying information about the external world- and this requires that information is present in sensory regions in a manner that is used by other brain regions^73^. The current results show that information about the world is reflected in brain activity patterns across a distributed set of brain regions, beyond low-level visual cortex, critically involving decision-related regions. It is debatable whether an additional requirement of a representational account of neural activity is that the encoded information is used by downstream neural systems in a manner that guides behavioural performance^73,74^. Processing of masked images merits additional consideration, particularly given the theoretical and experimental hurdles to isolate unconscious information processing^58^, and given that information that is not available for conscious report may not necessarily influence behaviour^13^. Pinpointing the specific brain signatures that determine whether or not unconscious contents influence behaviour awaits further determination.

Neurocognitive theories of consciousness propose that unconscious processing reflects feed-forward processing only, while local recurrent connections in sensory cortex are critical for bringing unconscious content into conscious awareness^75^. Visual signals that are embedded in noise and visually masked, are more likely to trigger feedforward processing only^76^. The neural network models used in our computer simulations of the visual task were all feedforward^41,77^, which may lack the capacity to preserve visual features across higher-order layers, so that any useful information might be left to local processes operating within each layer^78^. Therefore, in the presence of image perturbations (i.e. added Gaussian noise) the last readout layer of the FCNN may not fully exploit the information from previous layers to guide the perceptual decision. Likewise, in the human brain, unconscious feedforward processing may be able to produce information-rich representations in higher-order regions of the ventral visual pathway and even in parietal and prefrontal cortex, but without feed-back connections those representations are unable to guide behaviour and lead to conscious sensation. Recurrent feedback is thought to be critical for conscious experience^75,79,80^, though importantly, recent evidence indicates that long-range feedback connections from prefrontal cortex, rather than local feedback loops in visual cortex are more critical for visual consciousness^81^. The role of recurrent processing in unconscious information processing, however, remains unclear^82,83^, and there is suggestive evidence of a link between neural recurrency and unconscious processing too^84–86^. Recent modeling work indicates that recurrent neural networks (RNNs) may provide better representations than FCNN models^87,88^ and even explain brain activity better^78,88–90^. Although RNNs may be similar to deep FCNNs at least in the first pass, the evidence suggests that incorporation of feedback connections in RNNs can have an effect on the quality of the hidden representations^88,91^ and match performance with fewer layers relative to deep FCNNs^94^.

It will be relevant for future modeling work to investigate whether the addition of recurrent feedback connections to the FCNN model can improve the read-out of the hidden representations by the decision layer and hence improve classification performance of noisy images, or whether recurrent connections improve the informativeness of the hidden layer representation despite the FCNN classification performance remains at chance level with noisy images. We conclude that unconscious information processing in the visual domain, including processing without sensitivity, can lead to meaningful but hidden representational states that are ubiquitous in brains and biologically plausible models based on deep artificial neural networks. The work thus provides a framework for testing novel hypotheses regarding the scope of unconscious processes across different task domains.

## Methods

### Participants

Following informed consent, seven participants (mean = 29 years; SD = 2; 6 males) took part in return of monetary compensation at a rate of 20€ per each fMRI session. All of them had normal or corrected-to-normal vision and no history of psychiatric or neurological conditions. The study conformed to the Declaration of Helsinki and was approved by the BCBL Research Ethics Board. The present study used a high-precision, within-subject fMRI design similar to^92,93^, hence no statistical methods were used to predetermine sample sizes.

### Experimental procedure and stimuli

Subjects (N = 7) were presented with images of animate and inanimate items^43^. We selected 96 unique items (48 animate and 48 inanimate, i.e. cat, boat) for the experiment. These images could also be grouped by 10 subcategories (i.e. animal, vehicle) and 2 categories (i.e. living v.s. nonliving). The experiment was an event-related design. On each day, subjects carried out nine blocks of 32 trials each. Each block was composed of 16 animate and 16 inanimate items. Subjects performed six fMRI sessions of around 1 hour in separate days. There were hence 288 trials per day and 1728 trials in total per observer. The probe images were gray-scaled and presented in different orientations. The images were previously augmented using Tensorflow-Keras^95^. A random-phase noise background generated from the images was added to the target image before the experiment to facilitate masking.

The experiment was programmed using Psychopy v1.83.04^96^. The experiment was carried out on a monitor with a refresh rate of 100 Hz. A fixation point appeared for 500 ms, followed by a blank screen for 500 ms. Twenty frames of Gaussian noise masks were then presented and followed by the probe image, which was followed by another twenty frames of Gaussian noise masks. Then, there was a jittered blank period (1500 - 3500 ms) with a pseudo-exponential distribution in 500 ms steps (sixteen 1500 ms, eight 2000 ms, four 2500 ms, two 3000 ms, and two 3500 ms), selected randomly and without replacement on each block of 32 trials. Following the jittered blank period, participants were required (i) to identify the category of the image (ii) to rate the state of visual awareness associated with the image. There was a 1500 ms deadline for each response. For the categorization decision task, two choices were presented on the screen - living (V) and nonliving (nV) - i.e. ‘V nV’ or ‘nV V’ with the left-right order of the choices randomly selected for each trial. Subjects pressed ‘1’ (left) or ‘2’ (right) to indicate the probe condition. For the awareness decision task, there were 3 choices: (i) ‘I did not see anything that allowed me to categorize the item, I was completely guessing’; henceforth, the unconscious trials, (ii) partially unconscious (‘I saw a brief glimpse but I am not confident of the response’), and (iii) conscious (‘I saw the object clearly or almost clearly and I am confident of the categorization decision’). The intertrial interval then followed with a jittered blank period of 6000 - 8000 ms with a pseudo-exponential distribution in 500 ms steps. The asynchrony between probe images across successive trials therefore ranged between 11.5 and 15.5 seconds.

The duration of the probe image was based on an adaptive staircase that was running throughout the trials. Specifically, based on pilot tests, we elected to use an staircase to get a high proportion of unconscious trials while ensuring that perceptual sensitivity was not different from chance level. If the observer reported ‘glimpse’, the number of 10 ms frames of stimulus presentation was reduced by one frame for the next trial, unless it was already only one frame of presentation; if the observer reported ‘conscious’, the number of frames of presentation would be reduced by two or three frames for the next trial, unless it was less than two to three frames, in which case it would be reduced by one frame; if the observer reported ‘unconscious’, the number of frames increased by one or two frames, randomly, for the next trial. Examples of probe images are shown in Supplementary Figure 16

### Statistics

Non-parametric statistics were used for the ROC analyses. Samples were bootstrapped and the means of the bootstrapped samples were used for estimating the corresponding chance level. The reported p values were the probability of the chance level distribution being greater or equal to the mean of the samples. The statistics for these analyses were one-tailed because our study was designed to assess conditions in which the ROC was above the chance level. P values were corrected for multiple tests using Bonferroni, where relevant. All the analyses used non-parametric statistics except for two ANOVAs that are reported in the paper. For these parametric analyses, we verified the normality assumption using the Shapiro-Wilk test and the assumption of homogeneity of variances using the Levene test.

### Analysis of behavioral performance

We assessed whether the level of discrimination accuracy of the image departed from chance level in each of the awareness conditions. The metric to measure accuracy was A’, based on the area under the receiver operating curve (ROC-AUC)^45^. A response was defined as a ‘true positive’ (TP) when ‘living’ was both responded and presented. A response was defined as a ‘false positive’ (FP) when ‘living’ was responded while ‘nonliving’ was presented. A response was defined as a ‘false negative’ (FN) when ‘nonliving’ was responded while ‘living’ was presented. A response was defined as a ‘true negative’ (TN) when ‘nonliving’ was both responded and presented. Thus, a hit rate (H) was the ratio between TP and the sum of TP and FN, and a false alarm rate (F) was the ratio between FP and the sum of FP and TN. A’ was computed with different regularization based on 3 different conditions: 1) F ≤ 0.5 and H ≥ 0.5, 2) H ≥ F and H ≤ 0.5, 3) anything that were not the first two conditions. We first calculated A’ associated with the individual behavioral performance within each of the different states of awareness (henceforth called the experimental A’). Then, we applied permutation tests to estimate the empirical chance level. We bootstrapped trials for a given awareness state with replacement^97^; the order of the responses was shuffled while the order of the correct answers remained the same to estimate the empirical chance level. We calculated the A’ based on the shuffled responses and correct answers to estimate the chance level of the behavioral performance, and we called this the chance level A’.

This procedure was repeated 10,000 times to estimate the distribution of the empirical chance level A’ for each awareness state and each observer. The probability of empirical chance level A’ being greater or equal to the experimental A’ was the statistical significance level (one-tailed p-value,^98^). Hence, we determined whether the A’ of each individual was above chance across the different awareness states (Bonferroni corrected for multiple tests).

### Multivariate pattern analysis: decoding within each awareness state

Multivariate pattern analysis (MVPA) was conducted using Scikit-learn^104^ and Nilearn^105^ using a linear support vector machine (SVM) classifier. SVM has limited complexity, hence reducing the probability of over-fitting (model performs well in training data but bad in testing data) and it has been shown to perform well with fMRI data^106,107^. We used an SVM with L1 regularization, nested with invariant voxels removal and feature scaling between 0 and 1 as preprocessing steps. The nested preprocessing steps were fit in the training set and applied to the testing set. Note that these preprocessing steps are different from the detrending and z-scoring of the BOLD signals and represent conventional machine learning practices^108^.

During cross-validation, trials corresponding to one living (i.e. cat) and one non-living (i.e. boat) item for a given awareness state (i.e. unconscious) were left-out as the test set and the rest was used to fit the machine learning pipeline. With 96 unique items, 2256 cross-validation folds could be performed in principle. However, because the awareness states were randomly sampled for each unique item (i.e. cat), the proportion of examples for training and testing were not equal among different folds. Some subjects had less than 96 unique items for one or more than one of the awareness states. Thus, less than 2256 folds of cross-validations were performed in these cases. We elected to use the non-parametric ROC-AUC as a metric of classification performance given that it is robust to class imbalance and provides a sensitive and criterion-free measure of generalization^109^.

To get an empirical chance level of the decoding, the same cross-validation procedures were repeated by replacing the linear SVM classifier with a ‘dummy classifier’ as implemented in Scikit-learn, which makes predictions based on the distribution of the classes of the training set randomly without learning the relevant multivariate patterns. The same preprocessing steps were kept in the pipeline.

The mean difference between the true decoding scores and the chance level decoding scores was computed as the experimental score. To estimate the null distribution of the performance differences, we performed permutation tests. First, we concatenate the true decoding scores and the chance level decoding scores and then shuffle the concatenated vector. Second, we split the concatenated vector into a new ‘decoding scores’ vector and a new ‘chance level decoding scores’ vector. The mean differences between these two vectors were computed. This procedure was repeated 10,000 times to estimate the null distribution of the performance differences. The probability that the experimental score was greater or equal to the null distribution was the statistical significant level (one-tailed p-value, corrected for the number of ROIs using Bonferroni).

### Multivariate pattern analysis: generalization across awareness states

Here the classifier was trained from data in a particular awareness state (the ‘source’; e.g. on conscious trials) and then tested on a different awareness state (the ‘target’; e.g. on unconscious trials) on top of the cross-validation procedure described above. Similar to the decoding analysis within each awareness state, instances corresponding to one living and one non-living item in both ‘source’ and ‘target’ were left out, but only the left-out instances in the ‘target’ were used as the test set. The rest of the instances in ‘source’ were used as the training set to fit the machine learning pipeline (preprocessing + SVM) as described above. The performance of the fitted pipeline was estimated by comparing the predicted labels and the true labels using ROC-AUC for the test set.

To get an empirical chance level of the decoding, a similar procedure to that described above with a ‘dummy classifier’ was used here. Similar permutation test procedures were used to estimate the empirical null distribution of the difference between the experimental and chance level ROC-AUC and the estimation was repeated 10,000 times. The probability that the experimental score was greater or equal to the null distribution was the statistical significant level (one-tailed p-value).

### Computational model simulation

We used different pre-trained FCNN models (i.e. AlexNet, VGGNet, ResNet, MobileNet, and DenseNet) implemented in Pytorch V1.0^110^ to perform the simulations. These models were downloaded from PyTorch in December 2019. The FCNNs learned to perform the same visual discrimination task as the human observers. FCNNs were trained with clear images of animate and inanimate items. In order to make the network less sensitive to the noise added during testing, one example composed of random noise only was added to each batch of animate and inanimate images (batch size =8) during the training phase, with the noise sampled by a normal distribution with the mean and standard deviation of the images of the same batch.

The FCNNs were tested under different levels of noise in the image. The goal here was to emulate the pattern observed in the fMRI study (i.e. decoding of the noisy image in the absence of perceptual sensitivity) using a FCNN. To control for the initialization state of the FCNN models, we fine-tuned some of the popular FCNN pre-trained models, such as AlexNet^111^, VGGNet^112^, ResNet^113^, MobileNet^114^, and DenseNet^115^, which were pre-trained using the ImageNet dataset^116^ and then were adapted to our experiment using a transfer learning (fine-tuning) procedure^117^. After fine-tuning the FCNN models on the clear images used in the experiment, the models were tested on images with different noise levels.

A shallow RNN is equivalent to a deep ResNet with layers sharing weights among them. Directly implementing such a RNN leads to performance comparable to the deeper ResNet^94^. The AlexNet contained six convolutional layers; the DenseNet169 contained an initial convolutional layer and four consecutive dense blocks and each was followed by a transition convolutional layer, resulting in 168 convolutional layers; the MobileNetV2 contained an initial convolutional layer and seven bottleneck blocks, followed by a convolutional layer, a pooling layer, and another convolutional layer, resulting 25 convolutional layers; the Resnet50 contained an initial convolutional layer, followed by four convolutional blocks, resulting 50 convolutional layers; the VGG19 contained five convolutional blocks with increased size of convolutional processing, resulting 16 convolutional layers.

As shown in Supplementary Figure 17, pretrained FCNN models using ImageNet^116^ were stripped of the original fully-connected layer while weights and biases of the convolutional layers were frozen and not updated further^117^. An adaptive pooling^37^ operation was applied to the last convolutional layer so that the output of this layer became a one-dimensional vector, and a new fully-connected layer took the weighted sum of these outputs (i.e. the ‘hidden layer’). The number of artificial units used in the hidden layer could be any positive integer, but for simplicity, we took 300 as an example and we explored how the number of units (i.e. 2, 5, 10, 20, 50, 100, and 300) influenced the pattern of results. The number of hidden layer units determined the number of new weights, *w_i_*, for training. The outputs of the hidden layer were passed to an activation function^118^, which could be linear (i.e. identical function) or nonlinear (i.e. rectified function). A dropout was applied to the hidden layer during training but not during testing. Different dropout rates were explored (i.e. 0, 0.25, 0.5, and 0.75), where zero dropout rate meant no dropout was applied. The dropout operation was varied to investigate how feature representations could be affected by a simple regularization.

A new fully-connected layer, namely, the classification layer, took the outputs processed by the activation function of the hidden layer to compose the classification layer. The number of artificial units used in the classification layer depended on the activation function applied to the outputs of the layer. If the activation function was sigmoid (formula 1), one unit was used, while if the activation was a softmax function (formula 2), two units were used. Under subscripts ‘*i*’ denotes the *i^th^* output of a given artificial unit.

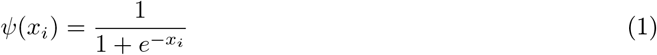

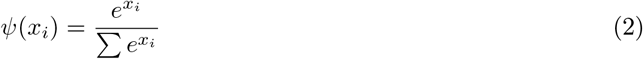

The re-organized FCNN was trained on the gray-scaled and augmented (flipped or rotated) experimental images and validated on images that were also gray-scaled but different degrees of augmentation. The loss function was binary cross-entropy. The optimizer was Adam^119^ with a learning rate of 1e-4 without decay. The validation performance was used to determine when to stop training, and the validation performance was estimated every 10 training epochs. The FCNN model was trained until the loss did not decrease for five validation estimations.

As noted, after training, the weights and biases of the FCNN model were frozen to prevent the model changing during the test phase. During the test phase, Gaussian noise was added to the images to reduce the FCNN classification performance. Similar augmentations as in the validation set were fed to the testing image sets. The noise added to the images was sampled from a Gaussian distribution centered at zero and different variance (σ). The level of noise was defined by setting up the variance at the beginning of each test phase.

The noise levels ranged from 0 to 1000 in steps of 50 following a logarithmic trend. For a given noise level, 20 sessions of 96 images with a batch size of 8 were fed to the FCNN model, and both the outputs of the hidden layer and the classification layer were recorded. The outputs of the classification layer were used to determine the ‘perceptual sensitivity’ of the FCNN model, and the outputs of the hidden layer were used to perform subsequent decoding analyses with a linear SVM classifier, in keeping with the fMRI analysis.

To determine the significance level of the FCNN model performance, the order of the true labels in each session was shuffled while the order of the predicted labels remained the same. The permuted performance was calculated for the 20 sessions. This procedure was repeated 10,000 times to estimate the empirical chance level of the FCNN model. The significance level was the probability that the performance of the FCNN model was greater or equal to the chance level performances (one-tailed test against 0.05). If the p-value is greater or equal to 0.05, we considered that FCNN performance was not different from the empirical chance level.

We then assessed, for a given noise in the image, whether the hidden layer of the FCNN (i.e. following the last convolutional layers), contained information that allowed decoding of the category of the image (living vs non-living). A linear SVM used the information contained in the FCNN hidden layer to decode the image class across different levels of noise, even when the FCNN model classification performance was at chance. The outputs and the labels of the hidden layer from the 20 sessions were concatenated. A random shuffle stratified cross-validation procedure was used in the decoding experiments with 50 folds to estimate the decoding performance of the SVM. The statistical significance of the decoding performance was estimated by a different permutation procedure to the FCNN, which here involved fitting the SVM model and testing the fitted SVM with 50-fold cross-validation in each iteration of permutation, and it was computational costly ^*b*^. On each permutation iteration, the order of the labels was shuffled while the order of the outputs of the hidden layer remained unchanged before fitting the SVM model^98^. The permutation iteration was repeated 100 times to estimate the empirical chance level. The significance level was the probability of the true decoding score greater or equal to the chance level. Because we were interested in those poorly performing FCNNs, we only attempted to decode the stimulus category from the hidden layer in those cases in which the FCNN classification performance was lower than 0.55 ROC-AUC.

### fMRI acquisition and preprocessing

A 3-Tesla SIEMENS’s Magnetom Prisma-fit scanner and a 64-channel head coil was used. In each fMRI session, a multiband gradient-echo echo-planar imaging sequence with an acceleration factor of 6, resolution of 2.4 x 2.4 x2.4 *mm*^3^, TR of 850 ms, TE of 35 ms, and bandwidth of 2582 Hz/Px was used to obtain 585 3D volumes of the whole brain (66 slices; FOV = 210mm). For each observer, one high-resolution T1-weighted structural image was also collected. The visual stimuli were projected on an MRI-compatible out-of-bore screen using a projector placed in the room adjacent to the MRI-room. A small mirror, mounted on the head coil, reflected the screen for presentation to the subjects. The head coil was also equipped with a microphone that enabled the subjects to communicate with the experimenters in between the scanning blocks.

The first 10 volumes of each block were discarded to ensure steady state magnetization; to remove non-brain tissue, brain extraction tool (BET,^99^) was used; volume realignment was performed using MCFLIRT^100^; minimal spatial smoothing was performed using a Gaussian kernel with FWHM of 3 mm. Next, Independent component analysis based automatic removal of motion artifacts (ICA-AROMA) was used to remove motion-induced signal variations^101^ and this was followed by a high-pass filter with a cutoff of 60 sec. The scans were aligned to a reference volume of the first session. All the processing of the fMRI scans were performed within the FSL (FMRIB Software Library; v6.0,^102^) framework and were executed using NiPype Python library^103^. Details of the NiPype preprocessing pipeline can be found in an online repository ^*c*^.

For each observer, the relevant time points or scans of the preprocessed fMRI data of each run were labeled with attributes such as (i.e. cat, boat), category (i.e. animal, vehicle), and condition (i.e. living vs. nonliving) using the behavioral data files generated by Psychopy (v1.84,^96^). Next, data from all sessions were stacked and each voxel’s time series was block-wise z-scored (normalized) and linear detrended. Finally, to account for the hemodynamic lag, examples were created for each trial by averaging the 3 or 4 volumes between the interval of 4 s and 7 s after image onset.

For a given awareness state, examples of BOLD activity patterns were collected for each of the 12 regions of interest (ROIs). There were 12 ROIs for each hemisphere (see Supplementary Figure 18). The ROIs included the lingual gyrus, pericalcarine cortex, lateral occipital cortex, fusiform gyrus, parahippocampal gyrus, inferior temporal lobe, inferior parietal lobe, precuneus, superior parietal gyrus, superior frontal gyrus, middle frontal gyrus, and inferior frontal gyrus (comprising pars opercularis gyrus, pars triangularis gyrus, and pars orbitalis gyrus). Automatic segmentation of the high-resolution structural scan was done with FreeSurfer’s automated algorithm recon-all (v6.0.0). The resulting masks were transformed to functional space using 7 degrees of freedom linear registrations implemented in FSL FLIRT^100^ and binarized. All further analyses were performed in native BOLD space within each observer.

## Acknowledgements

D.S. acknowledges support from the Basque Government through the BERC 2018-2021 program, from the Spanish Ministry of Economy and Competitiveness, through the ‘Severo Ochoa’ Programme for Centres/Units of Excellence in R & D (CEX2020-001010-S) and also from project grants PSI2016-76443-P and PID2019-105494GB-I00 from MINECO. R.S. acknowledges support by the Basque Government (IT1244-19 and ELKARTEK programs), and the Spanish Ministry of Economy and Competitiveness MINECO (project TIN2016-78365-R). The funders had no role in study design, data collection and analysis, decision to publish or preparation of the manuscript.

## Author Contributions Statement

N.M. and D.S. designed the study; N.M. analysed the data under the guidance of R.S. and D.S. N.M. prepared a first draft of the paper. All authors discussed the results and contributed towards the writing of the final version of the manuscript; D.S. supervised the project.

## Competing Interests Statement

The authors declare no competing interests.

## Data Availability

The fMRI data can be found at https://openneuro.org/datasets/ds003927

## Code Availability

Full scripts for the fMRI analyses and deep neural network simulations are available at https://github.com/nmningmei/unconfeats

## Supplementary Information

### Individual proportion of awareness ratings, target duration, probability of hits and false alarms and corresponding d’ and meta-d’ scores

To verify the pattern of results observed with A’, we also performed analyses of d’ and meta-d’. Hits were defined as responding non-living when a non-living object was presented and false alarms were defined as responding non-living when a living object presented. The d’ scores were computed across different awareness states within each subject. To compute d’, we first calculated the hit rate and the false alarm rate, and then converted the hit rate and the false alarm rate to the corresponding z scores using a standard normal distribution centered at zero with unit variance, with d’ being the difference between the hit rate z score and the false alarm z score. The meta-d’ was computed for each subject using all the awareness ratings, as a proxy of confidence, and measures the extent to which the subjective ratings track the correctness of the perceptual decision. We used a Python library, MetadPy^1^, to compute meta-d’ using the maximum likelihood estimate. The Supplementary Table 1 illustrates the proportion of each awareness state, the hit rate, the false alarm rate, d’, uncorrected p value of the d’, and the meta-d’ in Supplemental Table 1.

**Supplementary Table 1:** Individual proportion of awareness ratings, target duration, probability of hits and false alarms and corresponding d’ scores, as well meta-d’ scores for each subject.*:p < 0.05, **:p < 0.01, ***:p < 0.001; one-tailed p-values, Bonferroni corrected.

| Subject | Awareness | Proportion | Probe time (ms) | P(FA) | p(Hit) | d' | p | meta-d' |
| --- | --- | --- | --- | --- | --- | --- | --- | --- |
| Sub-01 | Unconscious | 0.45 | 18.77 ± 10.56 | 0.51 | 0.51 | 0.02 | 0.4371 | 2.53 |
|  | Glimpse | 0.26 | 29.96 ± 11.16 | 0.13 | 0.87 | 2.26 | < 0.001*** |  |
|  | Conscious | 0.29 | 41.30 ± 13.54 | 0.0 | 0.99 | 9.30 | < 0.001*** |  |
| Sub-02 | Unconscious | 0.35 | 16.48 ± 9.11 | 0.44 | 0.48 | 0.11 | 0.2341 | 1.64 |
|  | Glimpse | 0.45 | 24.22 ± 13.42 | 0.29 | 0.71 | 1.12 | < 0.001*** |  |
|  | Conscious | 0.2 | 36.74 ± 13.62 | 0.04 | 0.97 | 3.82 | < 0.001*** |  |
| Sub-03 | Unconscious | 0.44 | 34.73 ± 18.91 | 0.49 | 0.54 | 0.12 | 0.1675 | 1.54 |
|  | Glimpse | 0.31 | 50.79 ± 18.52 | 0.19 | 0.77 | 1.61 | < 0.001*** |  |
|  | Conscious | 0.24 | 58.92 ± 20.79 | 0.04 | 0.98 | 3.81 | < 0.001*** |  |
| Sub-04 | Unconscious | 0.44 | 34.92 ± 16.57 | 0.22 | 0.20 | -0.06 | 0.6526 | 1.85 |
|  | Glimpse | 0.32 | 51.55 ± 18.12 | 0.10 | 0.95 | 2.91 | < 0.001*** |  |
|  | Conscious | 0.24 | 59.07 ± 18.27 | 0.01 | 0.99 | 6.37 | < 0.001*** |  |
| Sub-05 | Unconscious | 0.42 | 19.80 ± 11.93 | 0.39 | 0.49 | 0.25 | 0.0276* | 2.55 |
|  | Glimpse | 0.28 | 31.43 ± 13.50 | 0.19 | 0.74 | 1.51 | < 0.001*** |  |
|  | Conscious | 0.29 | 40.22 ± 17.03 | 0.09 | 0.96 | 3.14 | < 0.001*** |  |
| Sub-06 | Unconscious | 0.44 | 22.70 ± 13.98 | 0.51 | 0.61 | 0.27 | 0.019* | 1.72 |
|  | Glimpse | 0.22 | 35.80 ± 15.42 | 0.21 | 0.77 | 1.53 | < 0.001*** |  |
|  | Conscious | 0.34 | 43.18 ± 17.83 | 0.05 | 0.91 | 3.00 | < 0.001*** |  |
| Sub-07 | Unconscious | 0.41 | 28.48 ± 16.39 | 0.39 | 0.57 | 0.47 | < 0.001*** | 1.37 |
|  | Glimpse | 0.39 | 41.19 ± 17.60 | 0.27 | 0.83 | 1.57 | < 0.001*** |  |
|  | Conscious | 0.20 | 50.47 ± 20.39 | 0.06 | 0.95 | 3.21 | < 0.001*** |  |

### Summary statistics of the MVPA results

**Supplementary Table 2:**
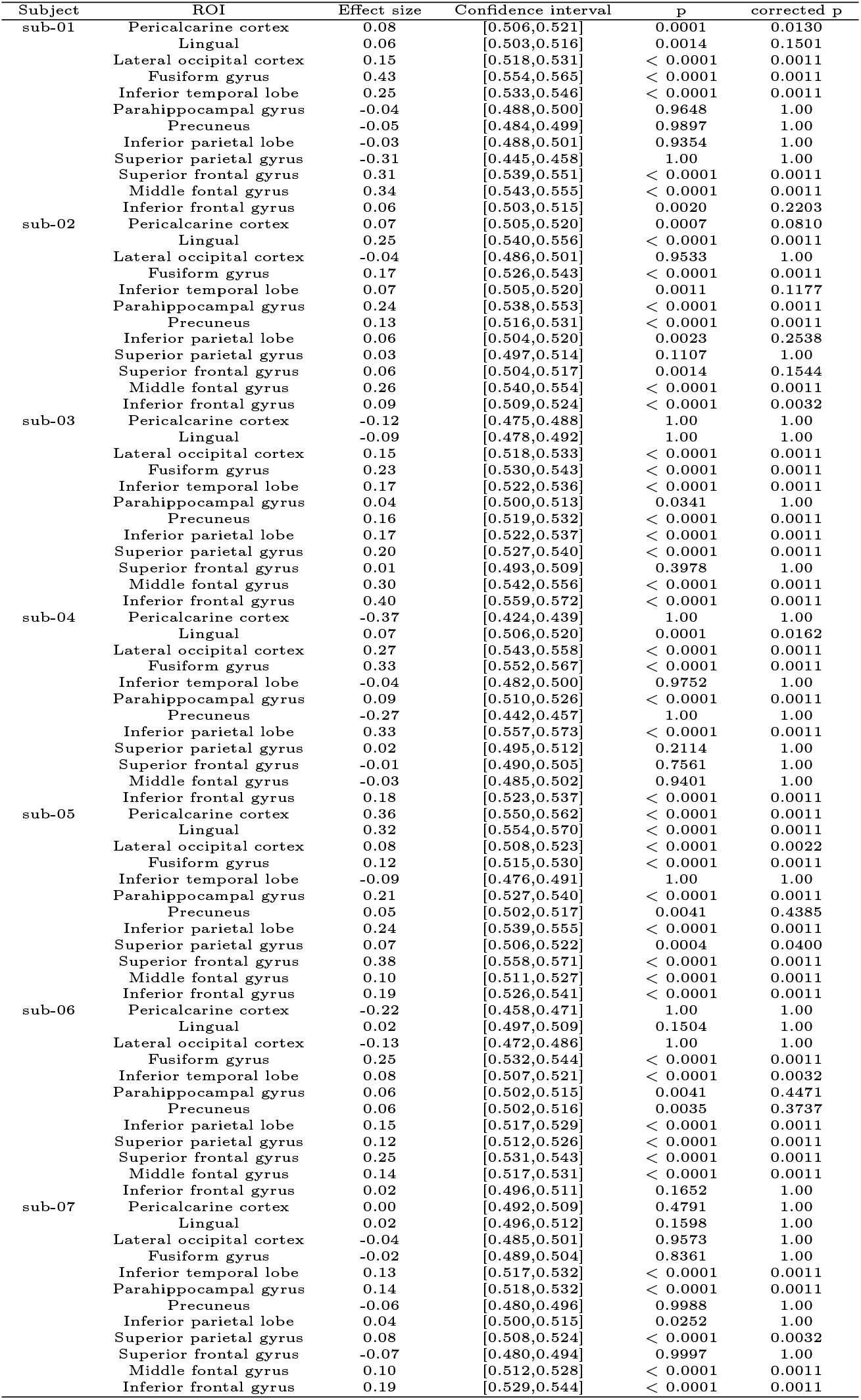
Summary statistics of the MVPA results of the unconscious trials. Cohen d was used as a measure of effect size ([mean ROC - chance ROC] / pooled s.t.d.). The confidence interval of the ROC scores were computed as follows: mean ROC +/- 1.96 * s.e.m.; s.t.d: standard deviation, s.e.m: standard error of the mean; one-tailed p-values, Bonferroni corrected.

**Supplementary Table 3:**
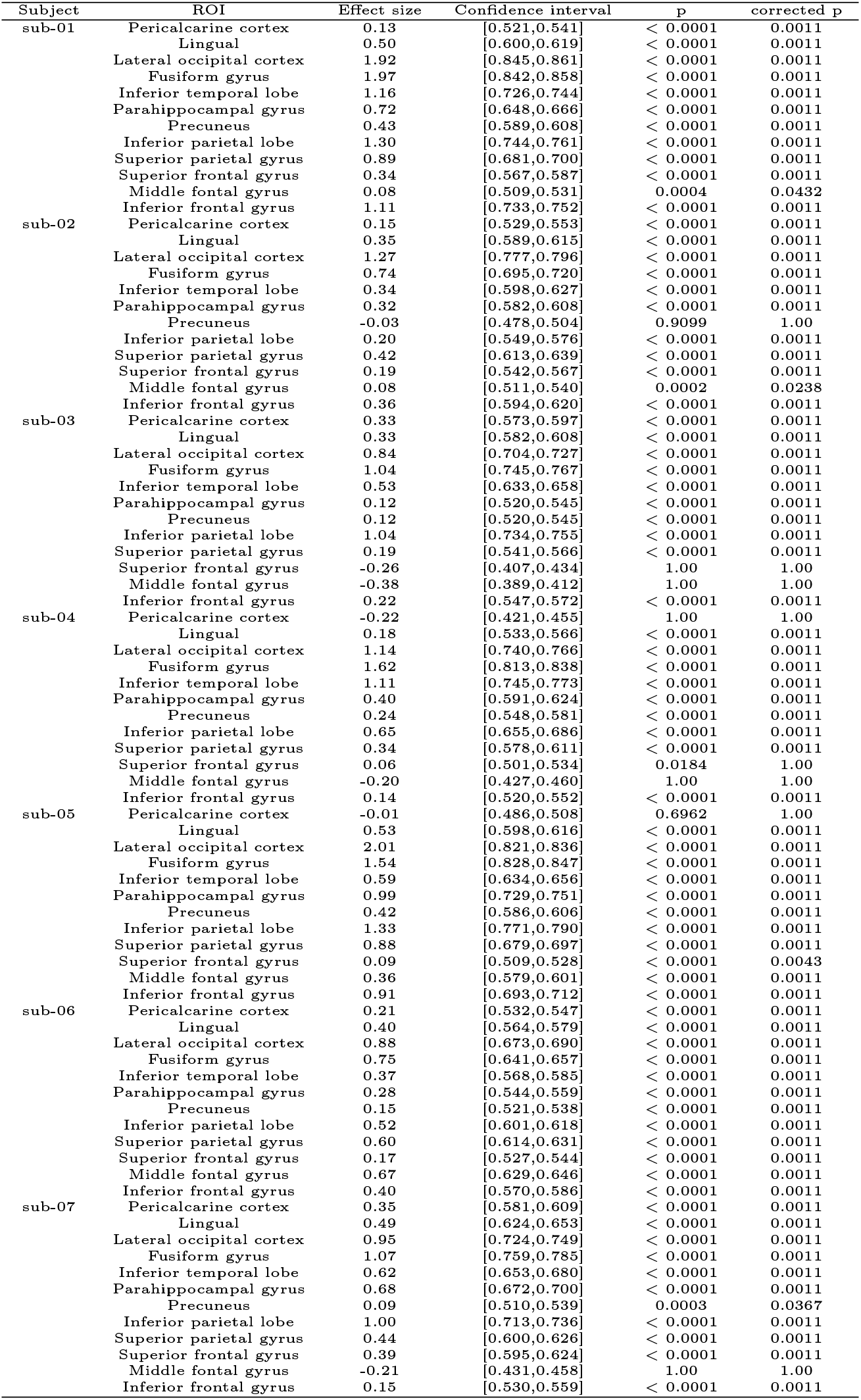
Summary statistics of the MVPA results of the conscious trials. Cohen d was used as a measure of effect size ([mean ROC - chance ROC] / pooled s.t.d.). The confidence interval of the ROC scores were computed as follows: mean ROC +/- 1.96 * s.e.m.; s.t.d: standard deviation, s.e.m: standard error of the mean; one-tailed p-values, Bonferroni corrected.

**Supplementary Table 4:**
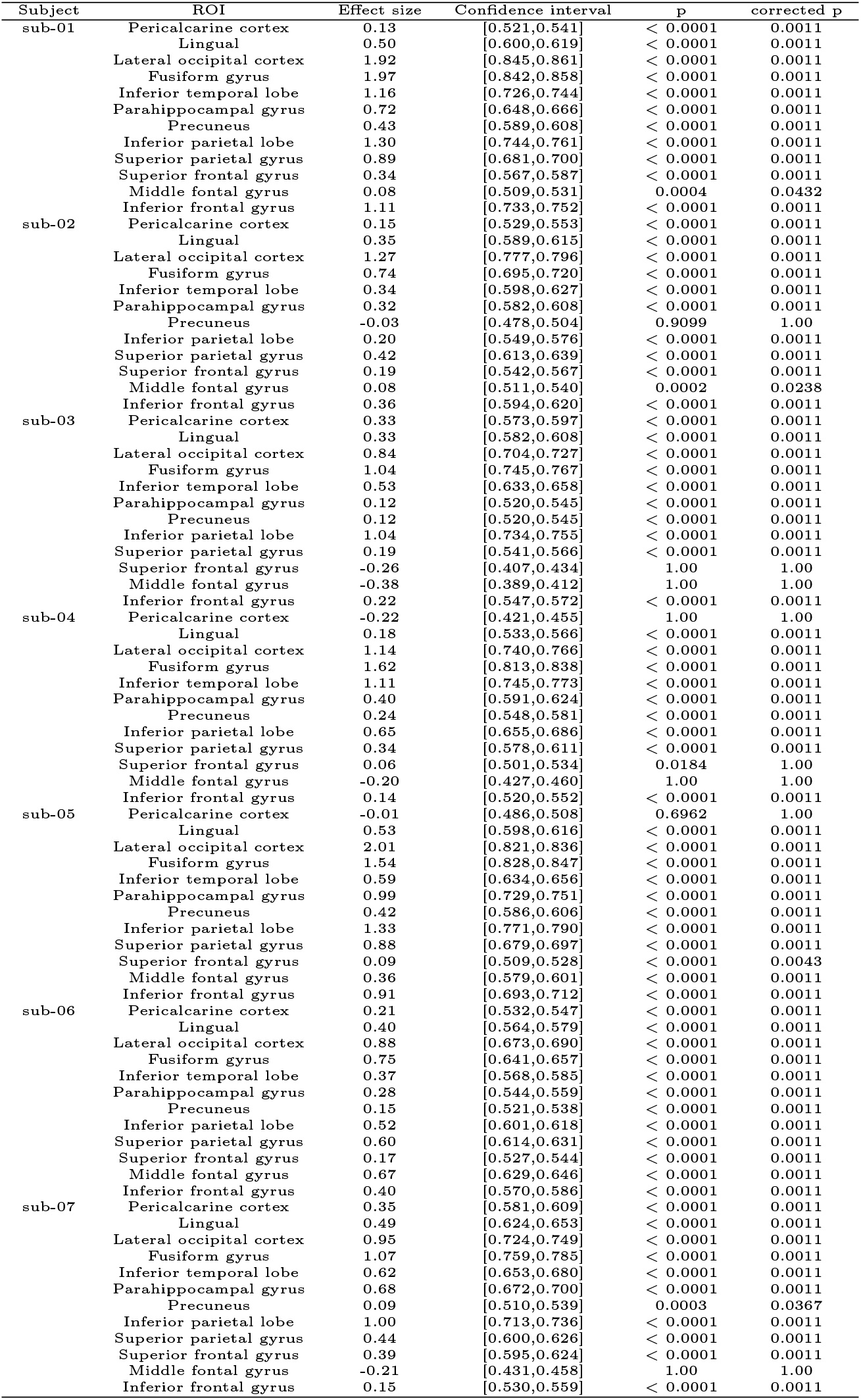
Summary statistics of the MVPA results of generalization from the conscious trials to the unconscious trials. Cohen d was used as a measure of effect size ([mean ROC - chance ROC] / pooled s.t.d.). The confidence interval of the ROC scores were computed as follows: mean ROC +/- 1.96 * s.e.m.; s.t.d: standard deviation, s.e.m: standard error of the mean; one-tailed p-values, Bonferroni corrected.

### Post-hoc tests of the feature importance

**Supplementary Table 5:**
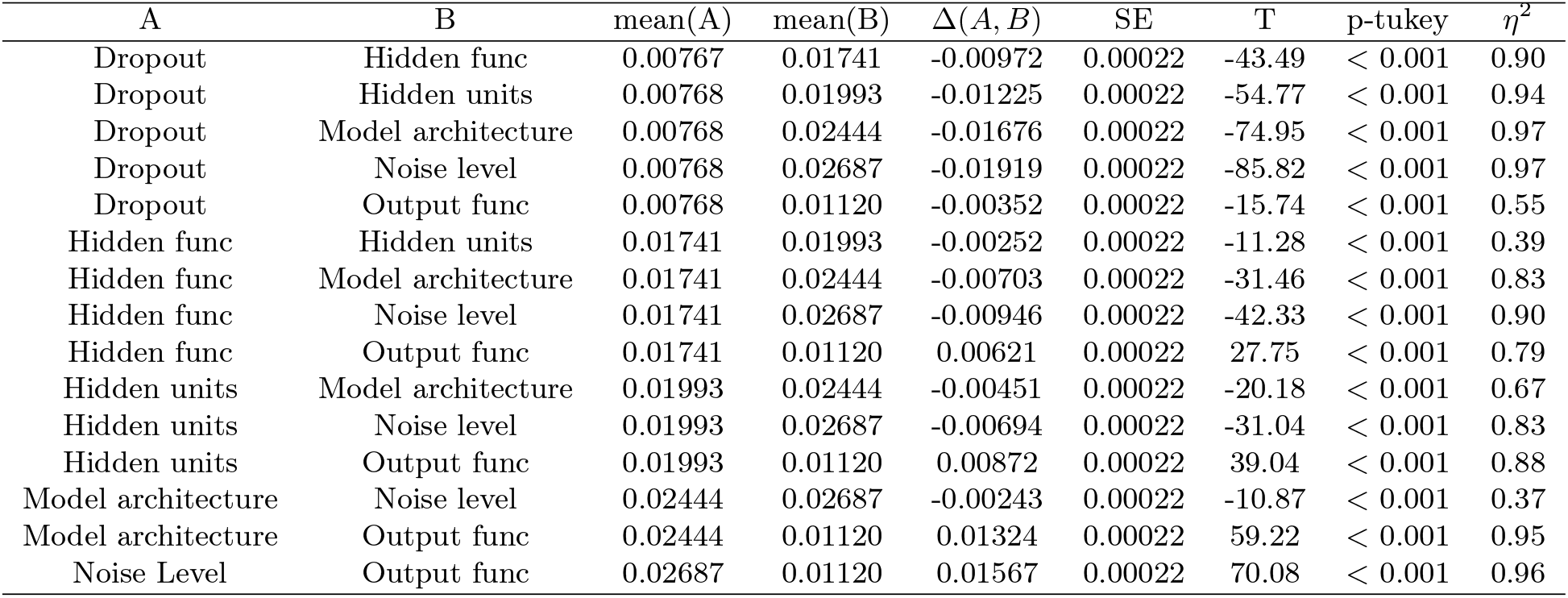
Post-hoc results regarding the feature importances. Dropout: Dropout rate; Hidden func: Hidden activation function; Output func: Output activation function.

### D’ scores across the different awareness states

**Supplementary Figure 1:**
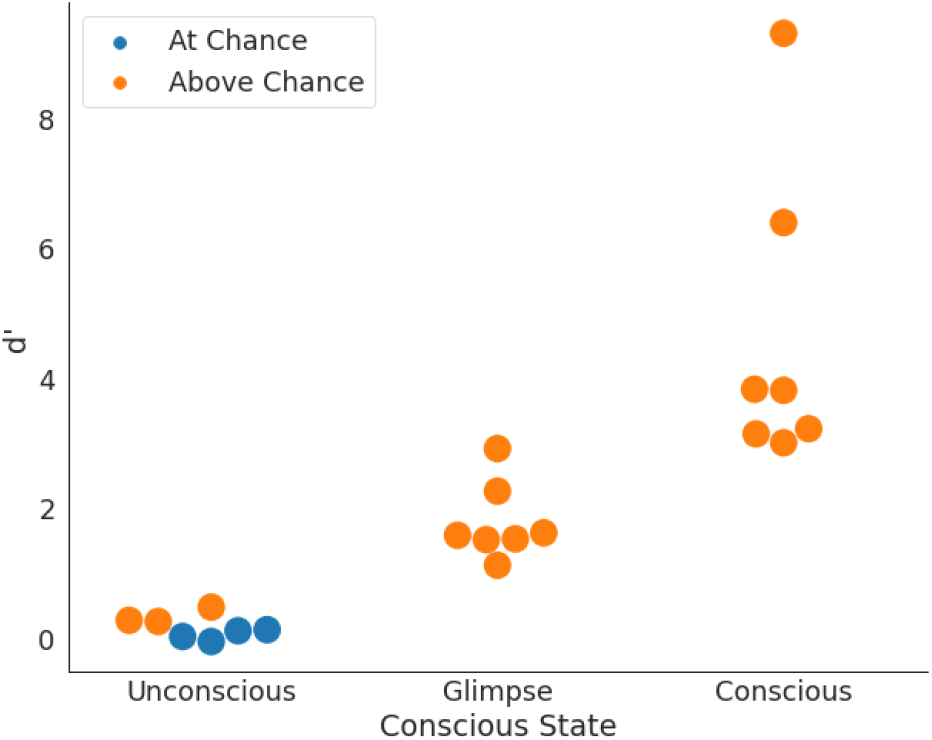
D’ scores across the different awareness states. We first calculated the hit rate and the false alarm rate, and then converted the hit rate and the false alarm rate to the corresponding z scores using a standard normal distribution centered at zero with unit variance. Permutation tests similar to those reported for A’ were run in order to assess -within each participant tested-, whether perceptual sensitivity differed from the empirical chance level. Blue dots depict the performance of those subjects that differed from the empirical chance level (uncorrected p values for the number of tests done across each of the awareness states). Orange dots depict the performance of those subjects that remained within the empirical chance level.

### Decoding performance

Across the different ROIs, we pooled the decoding accuracy of the four participants that displayed null perceptual sensitivity on trials rated as unaware and likewise for those participants that displayed above chance sensitivity. The Figure S2 illustrates these data. There are no apparent and consistent differences across the different ROIs.

**Supplementary Figure 2:**
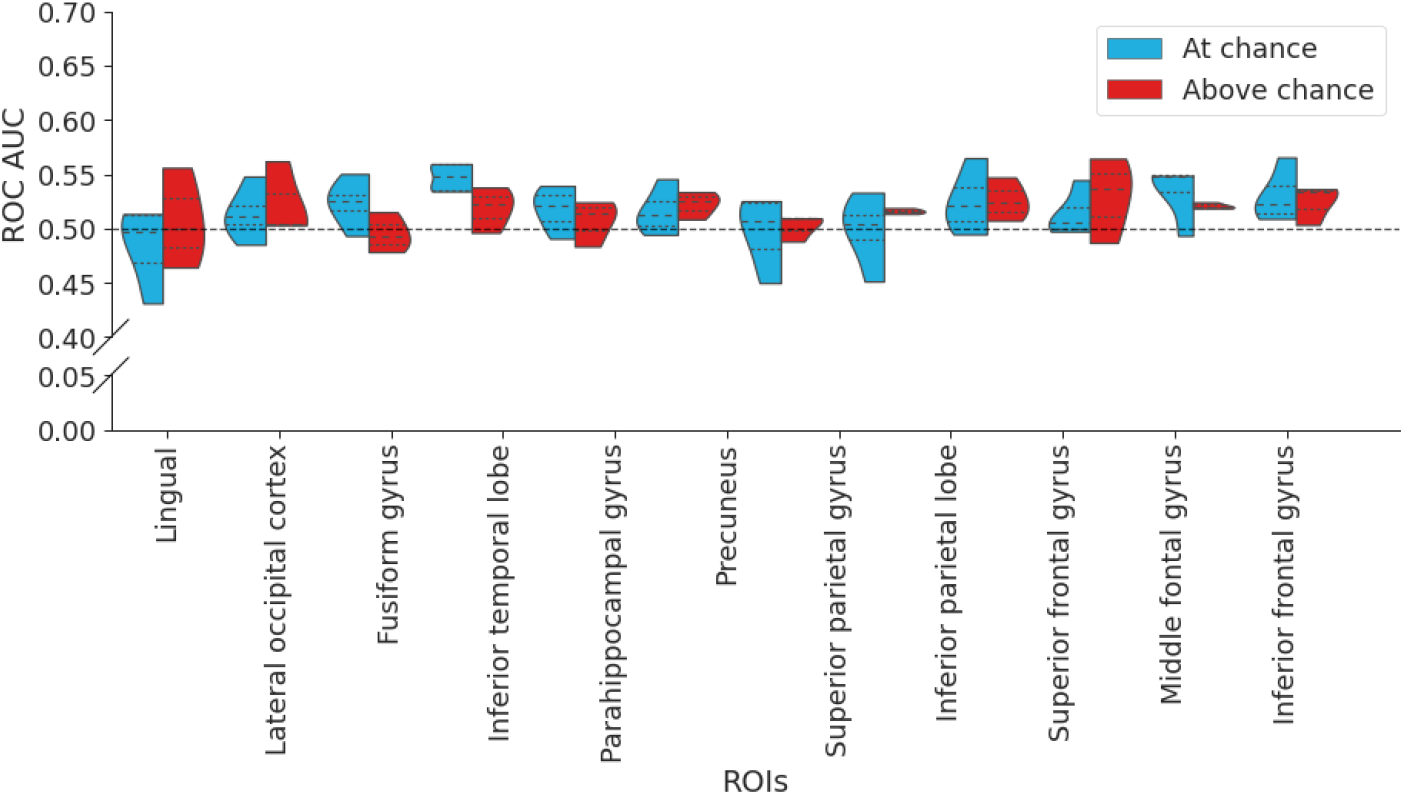
Distribution of decoding accuracy across the participants whose perceptual sensitivity was at chance and those who deviated from chance, including the mean, first and third quartiles.

### Additional MVPA analyses

**Supplementary Figure 3:**
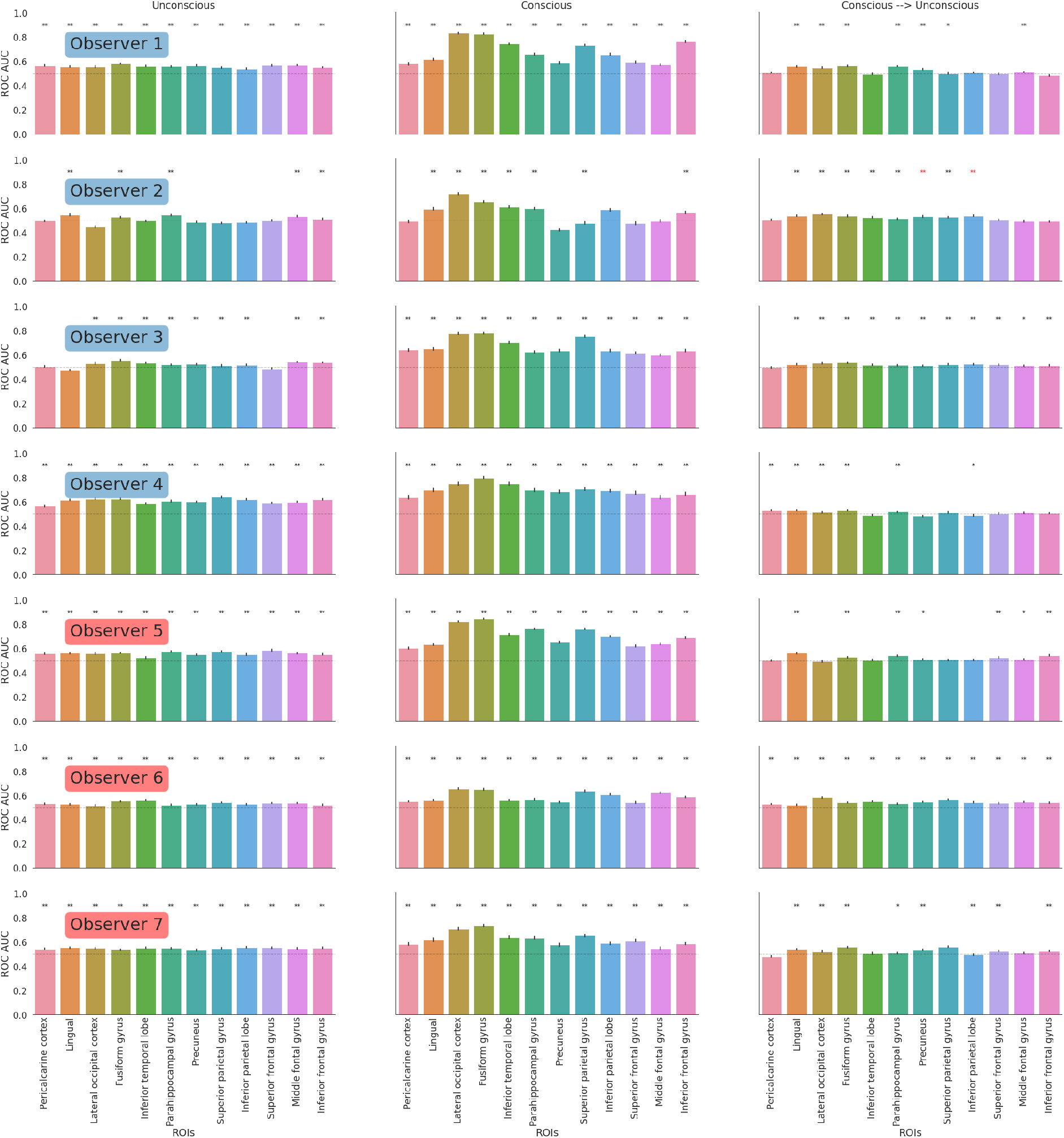
Decoding performance of the SVM with L1 regularization (C = 5) using the previous out-of-sample generalization cross-validation partitioning scheme: *:p < 0.05, **:p < 0.01, ***:p < 0.001; one-tailed p-value, after multiple comparison correction for the number of ROIs tested for each observer. Error bars represent the standard error of the mean. Red asterisks indicate those ROIs in which the cross-awareness state generalization appeared above chance but the decoding in the conscious condition was at chance, and these were discarded.

**Supplementary Figure 4:**
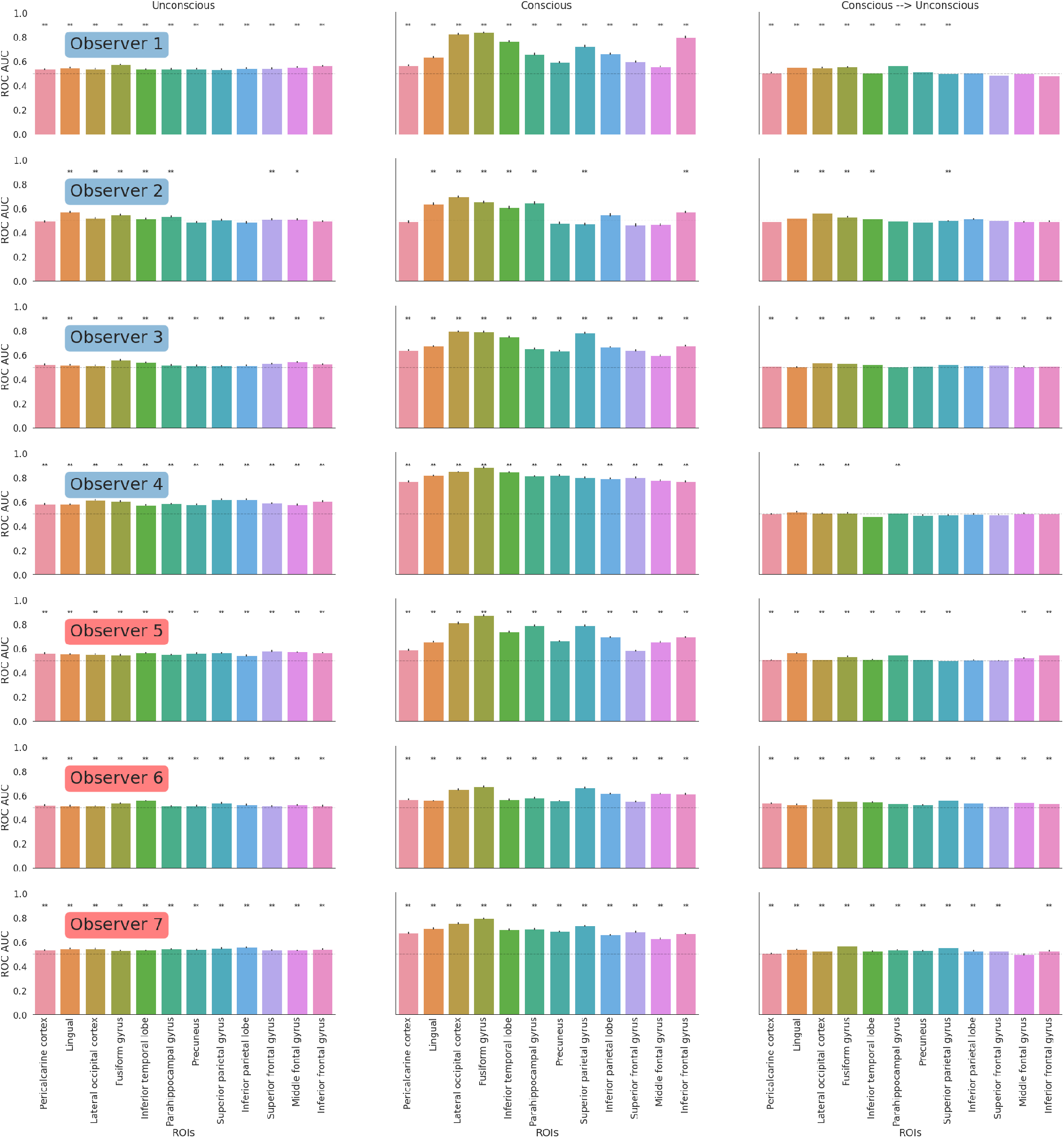
Decoding performance of the SVM with the default L1 regularization (C = 1) using the stratified random shuffle cross-validation partitioning scheme. During each cross-validation within each state of awareness, 95% of the data was used for training and 5% of the data was used for testing. This cross-validation scheme was repeated 1000 times. During the cross-awareness crossvalidation, 95% of the trials in the conscious condition were randomly selected for training and 100% of the trials in the unconscious condition were used for testing: *:p < 0.05, **:p < 0.01, ***:p < 0.001; one-tailed p-value, after multiple comparison correction for the number of ROIs tested for each observer. Error bars represent the standard error of the mean.

### Feature importance

**Supplementary Figure 5:**
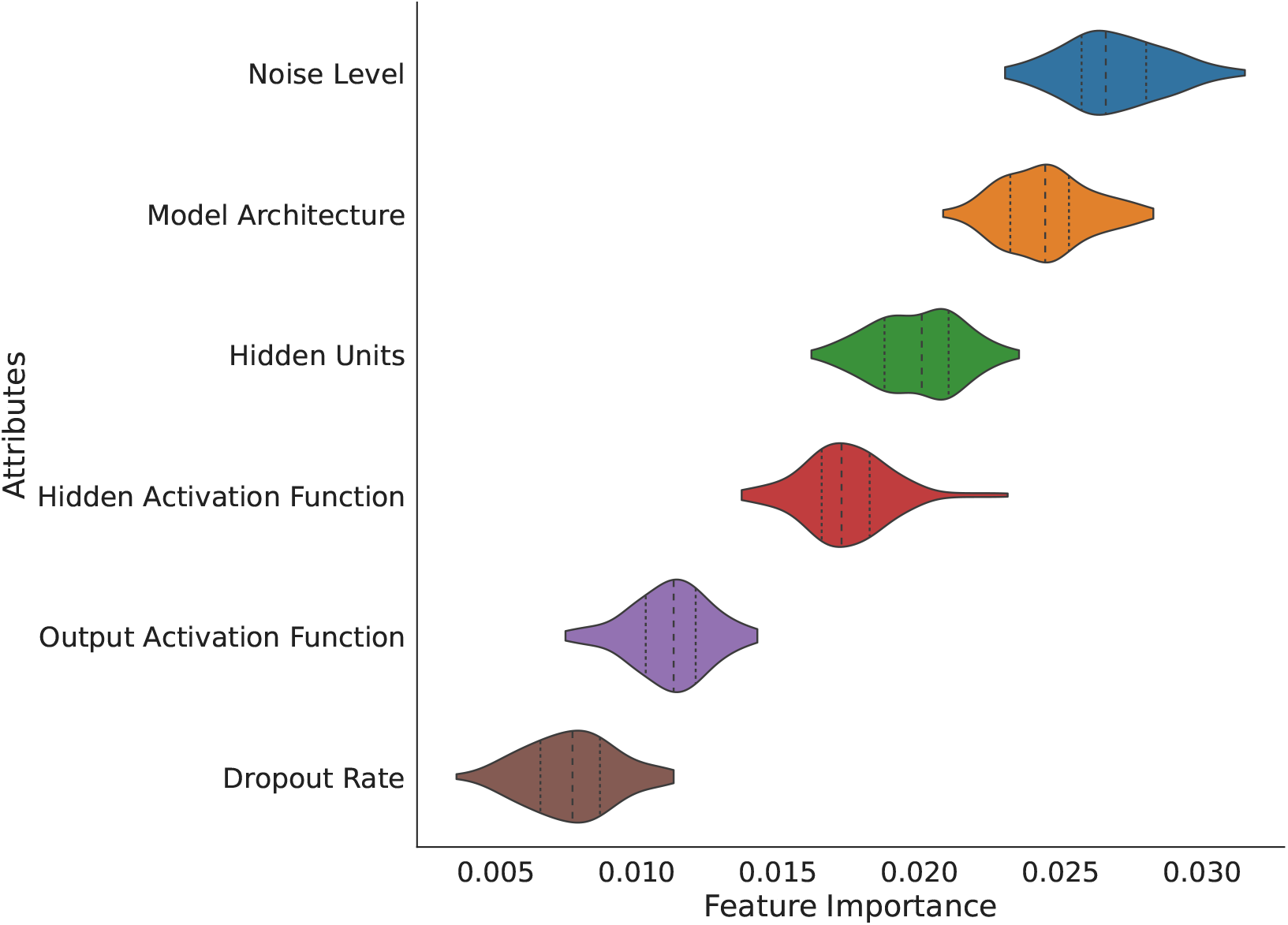
Feature importance of FCNN components that were manipulated in computational modeling. Feature importance was measured in arbitrary units, including the mean, first and third quartiles. The number of hidden layer units, noise levels, and pre-trained configurations influenced the decoding performance of the image class based on the hidden layer of the FCNN when its classification performance was at chance.

### Decoding hidden representations using clear and noisy images

Due to computational costs and limited resources, we run the simulation including the clear images using two FCNN models, the VGG19 and Resnet50. These two models were selected because they are known to predict a relatively high amount of variance in brain activity during visual processing^2^ (Brain-score board was checked on June 2021). The pipeline was otherwise similar to the original simulations.

**Supplementary Figure 6:**
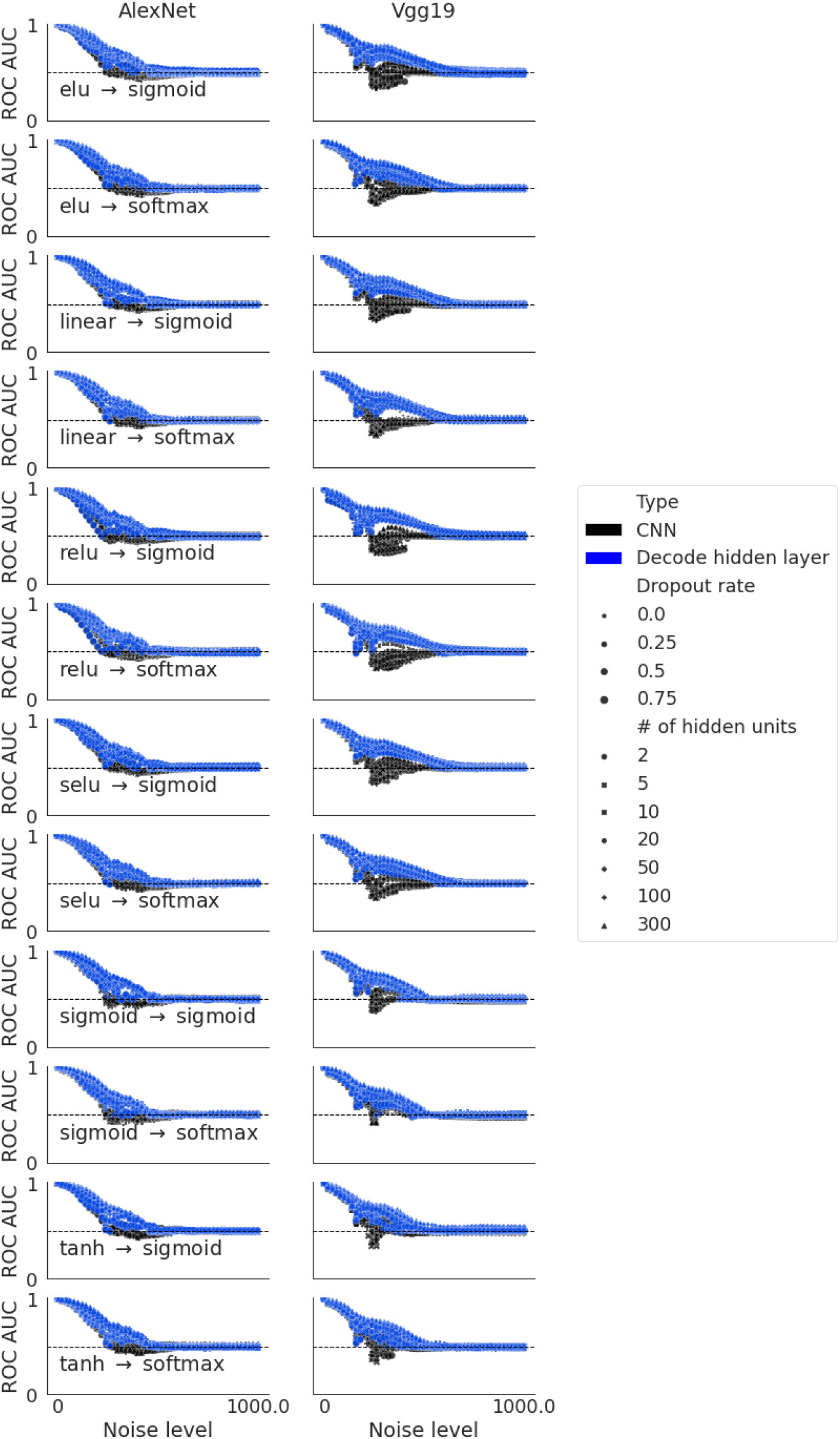
The black dots illustrate the classification performance of the FCNNs (VGGNet on the left and ResNet on the right) as a function of noise level. The blue dots illustrate the classification performance of the linear SVMs applied to the hidden layer of the FCNN models.

We also computed the performance difference between the FCNN and the SVM applied to the hidden layer and we can see in the Supplementary Figure 7 that there is a range of noise where the hidden layer contains more information that is decodable.

**Supplementary Figure 7:**
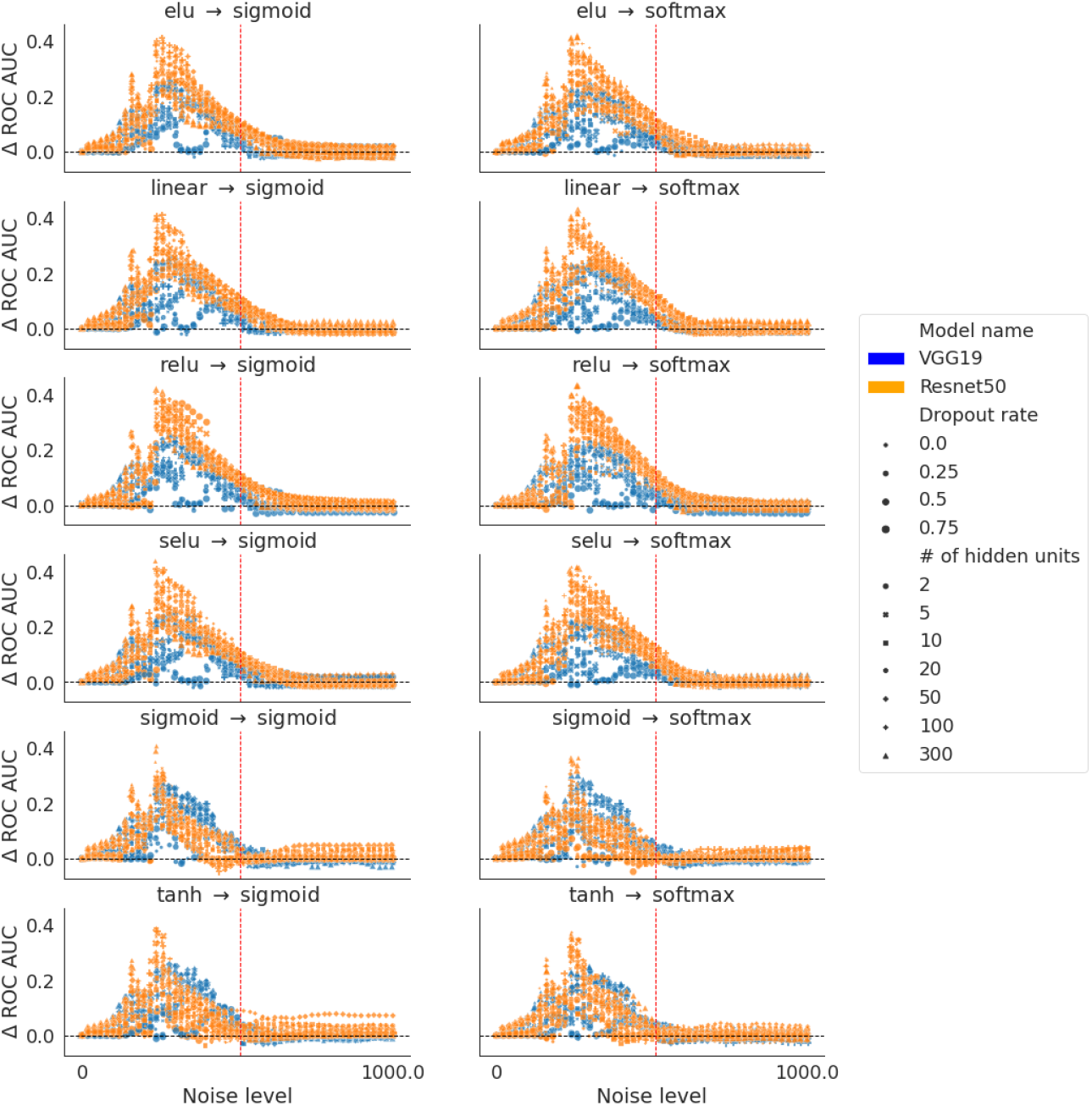
Difference in classification performance between the FCNN and the SVM applied to the hidden layer.

### Decoding the hidden layer representations when the FCNNs were trained with images embedded in noise

**Supplementary Figure 8:**
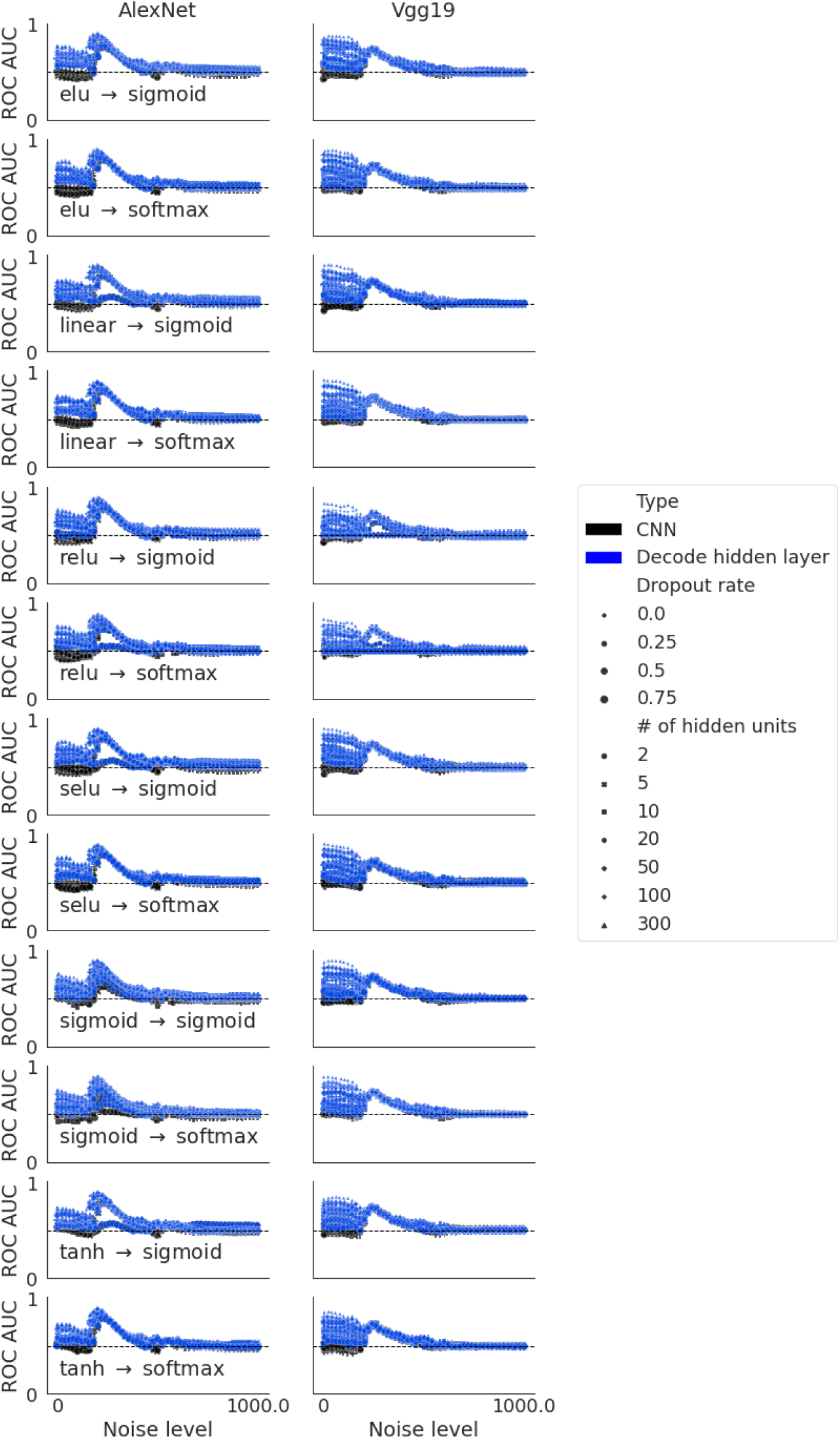
Performance of the FCNNs (VGG19 and Resnet50 with different configurations) and the SVM decoding the hidden representations, when the FCNNs were trained with images embedded in standard Gaussian noise. Color blue depicts the ROC AUC scores of the FCNNs and color black depicts the ROC AUC scores of the SVMs. Different sizes depict the different dropout rates and different shapes depict different numbers of hidden units.

**Supplementary Figure 9:**
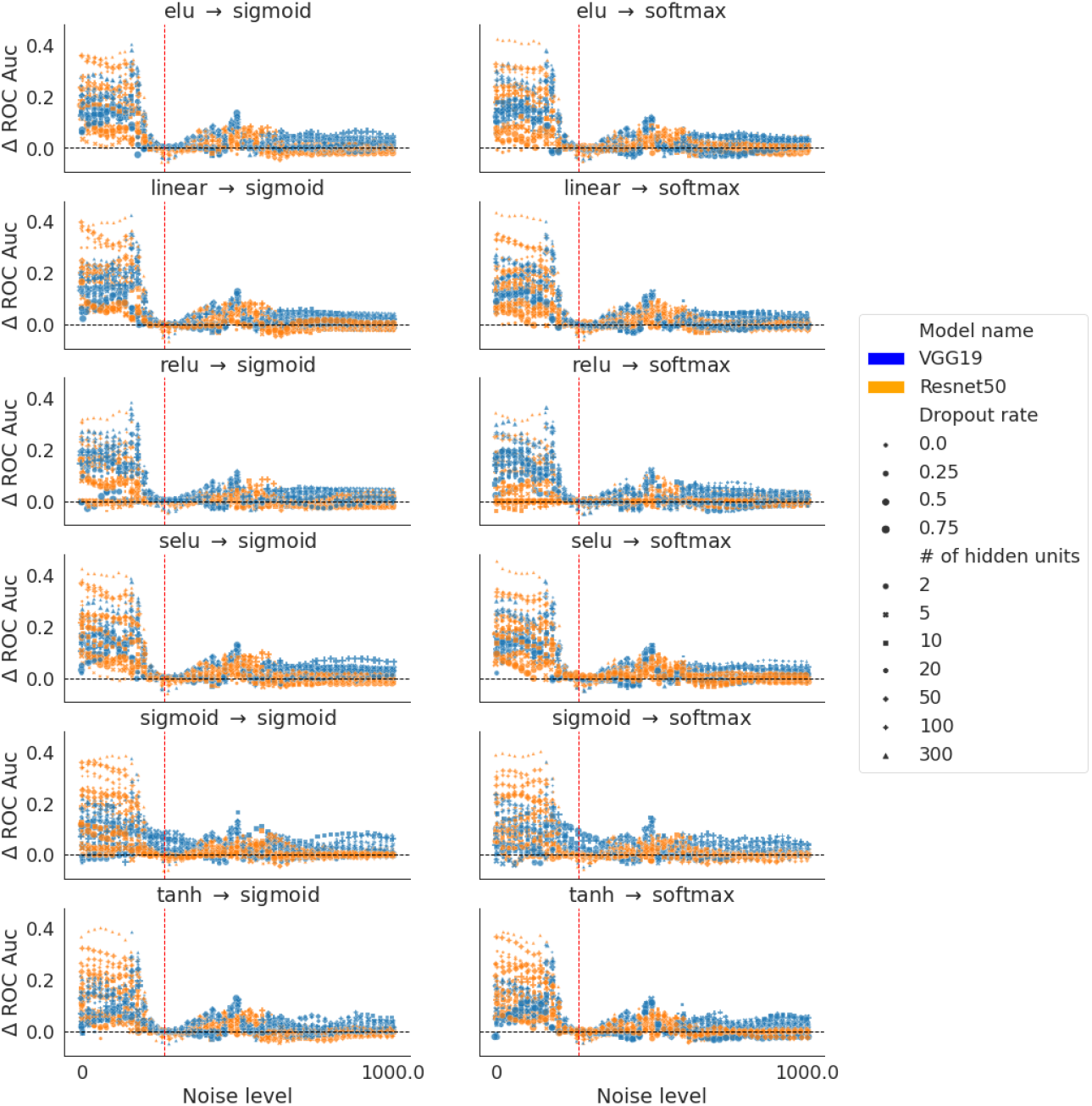
Difference of performance between the FCNNs and the SVM decoding the hidden layer representation..

**Supplementary Figure 10:**
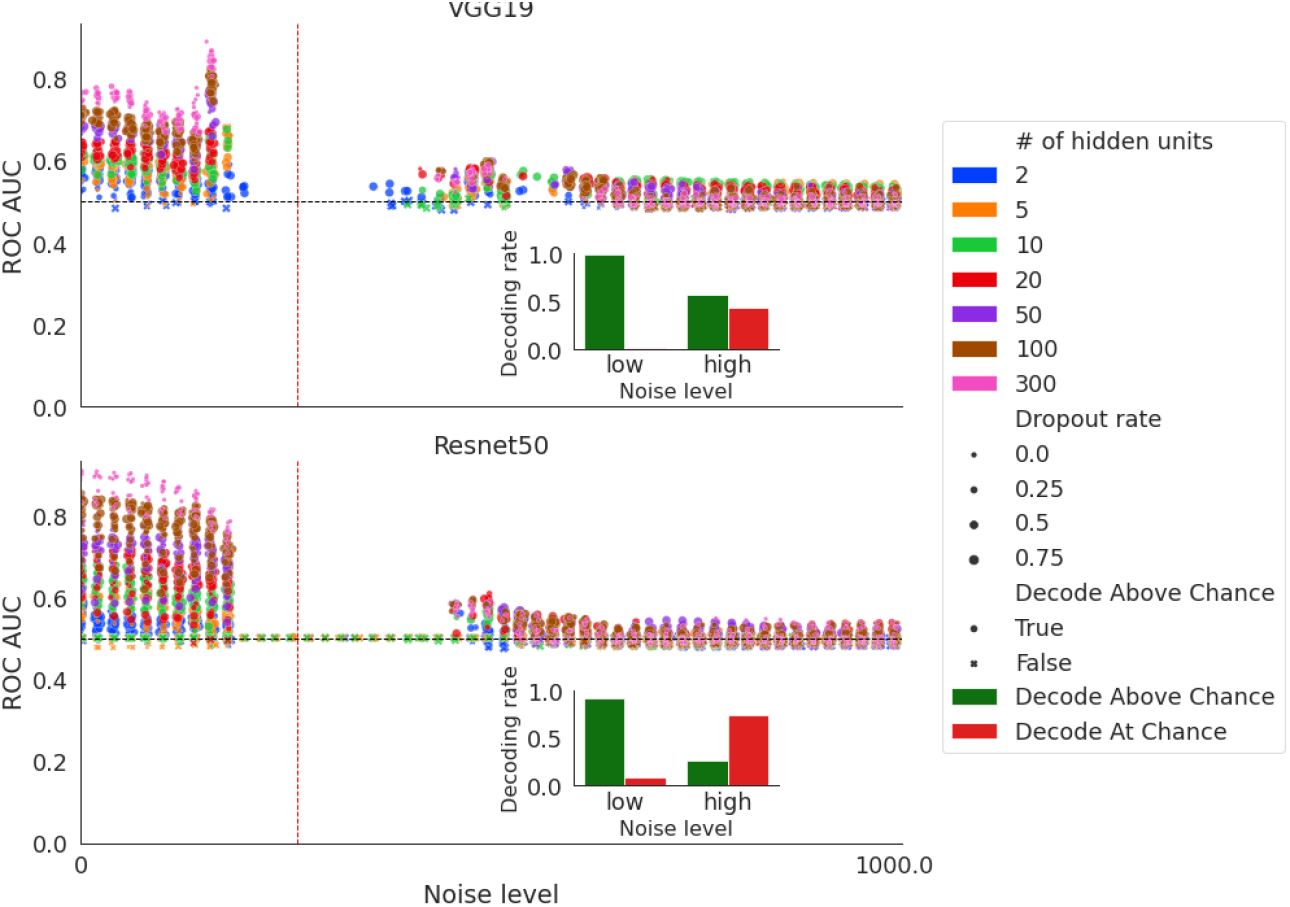
The performance of the SVM decoding the hidden representations when the FCNNs performance was at chance. The vertical red dash line depicts the noise level that was used during training. Low and high noise levels were defined based on this. The smaller figure within each subplot shows the proportion of FCNNs configurations that produced informative hidden representations when the noise level was low or high.

### Decoding first layer representations using clear and noisy images

Additional analyses were conducted to test whether the image categories could be decoded from the first layer activity pattern. Since the convolutional layers were frozen during the simulation, testing the decoding of the image categories from the first layer was redundant across different FCNN configurations (e.g. across different numbers of hidden units or activation functions). For the analysis of the first layer, we chose the two most studied CNN models, the VGG19 and Resnet50,, and simply added a new output layer on top of the original model that was trained on the ImageNet dataset. We trained the newly added output layer, which mapped 1000 outputs (ImageNet dataset categories) to 2 outputs (animate v.s. inanimate) on the clear images. The training only affected the weights associated with the last layer of the original CNN models and the newly added output layer. The output layer activation function was softmax and the loss function for training was the binary cross entropy. After the training, we froze the model and then, this was tested with images of different levels of noise following the pipeline of the first simulations. During the testing, the activity patterns of the first layer were recorded for further analyses. The image sizes were 128 x 128 x 3 and the output of the first layer for the VGG19 was 127 x 127 x 64 (i.e. with a dimensionality of over one million features after flattening). Relative to the number of trials (i.e. 96 x 20 =1920), the decoding matrix for a linear SVM classifier with L2 regularization implemented by Scikit-learn was computational intensive. In a unit test, it took the classifier 20 minutes to train on 80% of data and test on 20% of the data. For a standard 50-folds cross-validation, it took about 16 hours. To reduce the computational cost, we applied a principal components analysis (PCA) implemented by Scikit-learn to reduce dimensionality of the data. The PCA algorithm was set to keep the components that represented 90% of the variance. Then a linear SVM classifier with L2 regularization was applied to decode the image categories (animate and inanimate) in a 50-fold cross-validation. The discrimination performance of the FCNN and the SVM decoding performance based on the first layer activity patterns were compared to their corresponding empirical chance levels to assess the statistical significance using the same non-parametric t-test conducted for the analyses of the hidden layer. The results showed that the image categories could be decoded from the first layer and decoding performance decreased with increasing levels of noise in the image. Remarkably, even when the FCNN performed at chance level with noisy images, the image category could be decoded from the first layer activity patterns. These results are presented in Supplementary Figure 11.

**Supplementary Figure 11:**
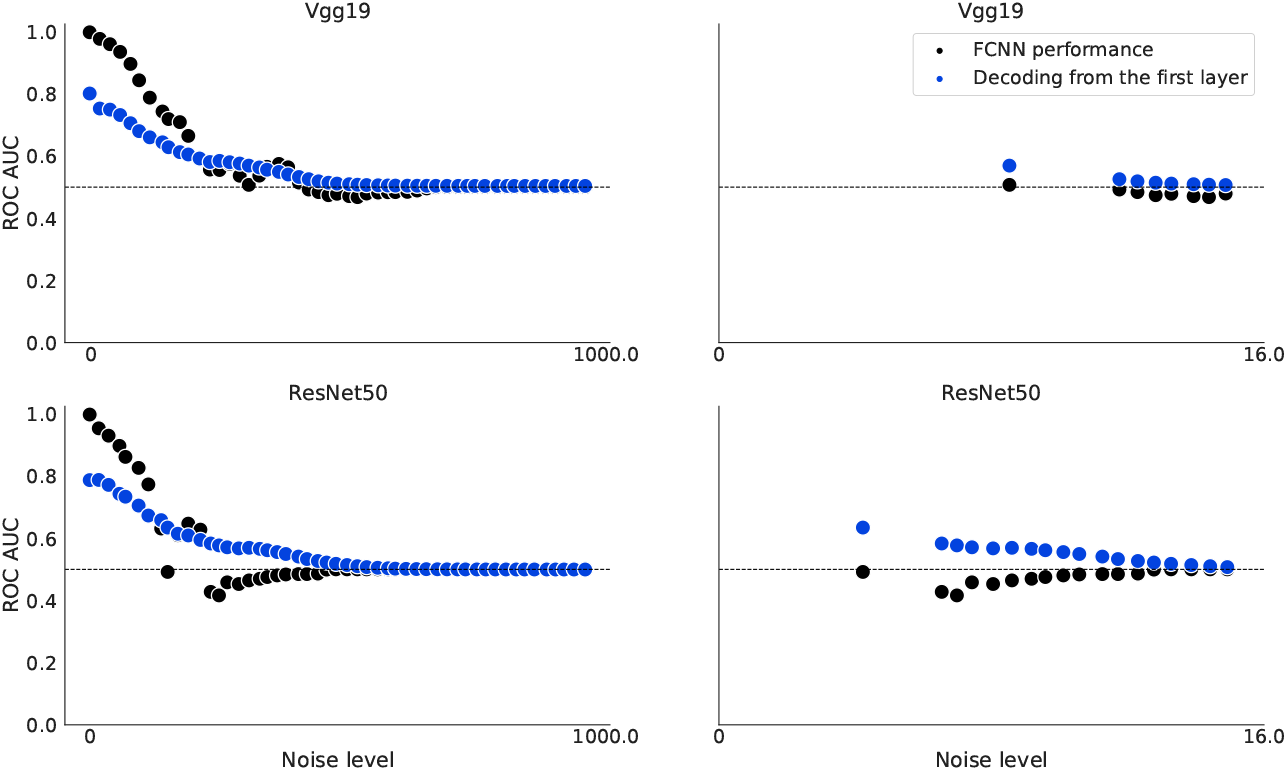
The left part depicts the FCNN performance (black dots) relative to the SVM decoding performance based on the first layer activity patterns (blue dots). The right part illustrates the range of noise at which the FCNN is at chance but the SVM could decode significantly above chance the image categories.

### Representational similarity analysis (RSA)

We used the VGG19 and the Resnet50 for the RSA because, as noted above, they are known to predict human brain responses better than other standard CNN models^2^. First, we fine tuned the pretrained models with the Caltech101 dataset to control for the size of the hidden representations (i.e. to match the number of units across models for the RSA). An adaptive pooling was applied to the last convolutional layer of the FCNN models so that the outputs became 1-dimensional vectors. The pooled activity patterns of the VGG19 had 512 units while the activity patterns of the Resnet50 had 2048 units. Then, a fully-connected layer with 300 units was added to the outputs of adaptive pooling, followed by a scaled exponential linear unit function (SELU^3^). SELU was chosen to speed up the train of the new components and scale the output activity patterns for better propagating information to the next layer.

A fully-connected layer with 96 units (1 per category of the Caltech dataset) was added to the outputs of the SELU layer, followed by a softmax activation function to compute probabilistic predictions of the image categories. The model was trained using 96 unique categories of the Caltech101 images (BACK-GROUND_Google, Faces, Faces_easy, stop_sign, and yin_yang were excluded,^4^). The convolutional layers were frozen during the training. The loss function was binary cross entropy and the optimizer was stochastic gradient descent. The data was split into train and validation partitions and the training was terminated if the performance on the validation data did not improve for five consecutive epochs.

We then fed the trained FCNN models with exactly the same images used in our experiment but without the noise background, in order to extract the hidden representations of the images. We then average the hidden representations of the trials belonging to the same item (i.e cat) and computed the model RDM for both the VGGNet and the ResNet (Supplementary Figures 12 and 13 depict the RDM for these hidden representations and it can seen there are clear clusters for the image categories). We then extracted the BOLD activity patterns using a mask that contained all the ROIs used for the decoding analyses. We averaged the BOLD signals of trials belonging to the same item (i.e cat) within a searchlight sphere of 6 mm that moved around the brain mask. A representational dissimilarity matrix (RDM) was computed for the extracted BOLD signals using Pearson correlation among each pair of images within each sphere. The lower triangles of the two RDMs were extracted and correlated using 1 - Spearman correlation for each sphere of the searchlight. The center of the sphere was assigned the Spearman correlation coefficient. These results are however descriptive, since currently there are no robust within-subject RSA analytical pipelines to make within-subject statistical inference.

The resulting brain map of FCNN model/brain similarities was converted to standard space for visualization using Nipype and FSL tools. This pipeline was run separately in the unconscious and conscious conditions, separately within each participant. The results are depicted in Supplementary Figures 14 and 15 (for the unconscious and conscious RSA analyses respectively).

**Supplementary Figure 12:**
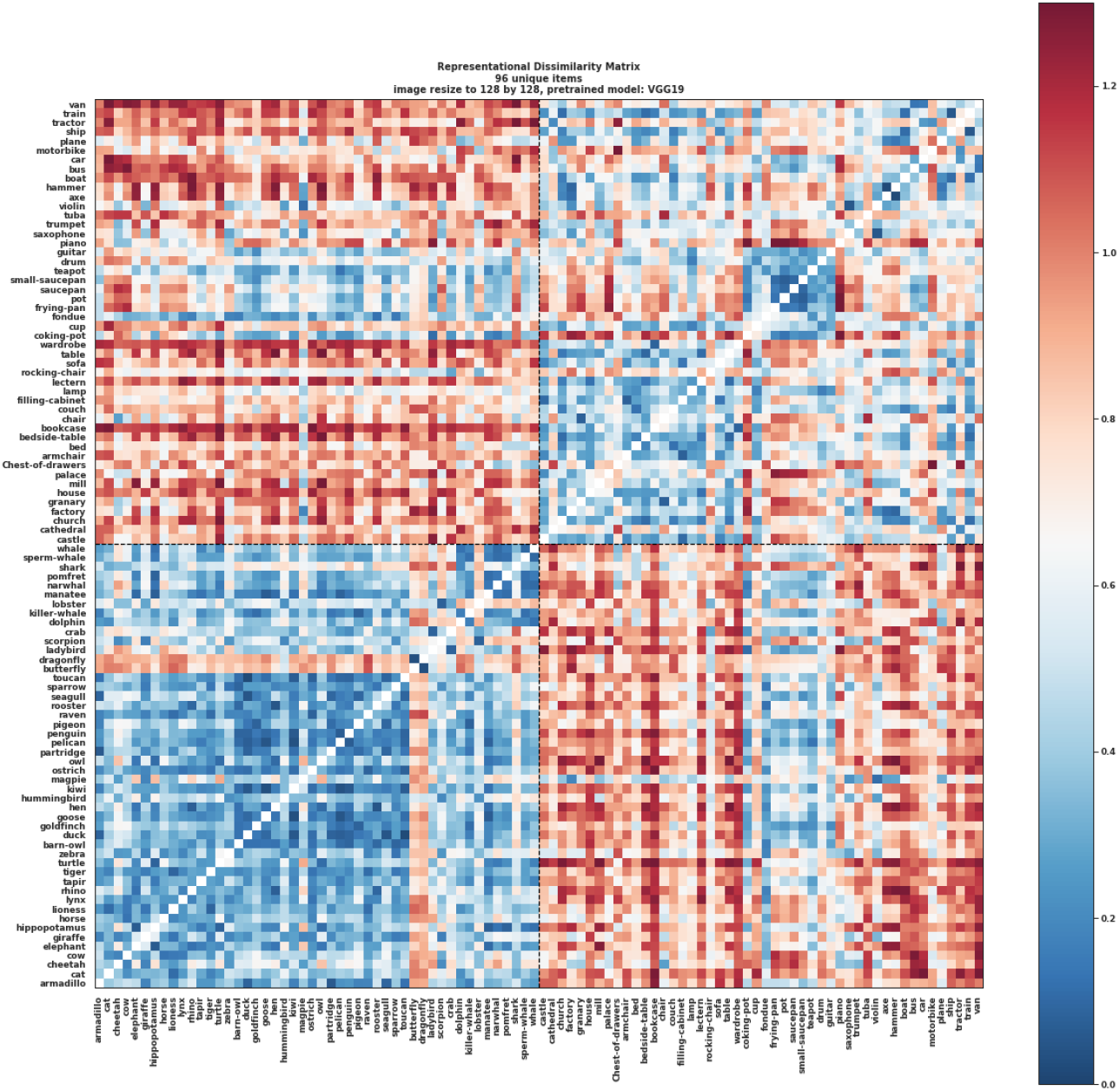
Representational dissimilarity matrix of the hidden representations of the VGG19 model fined-tuned by Caltech101 dataset.

**Supplementary Figure 13:**
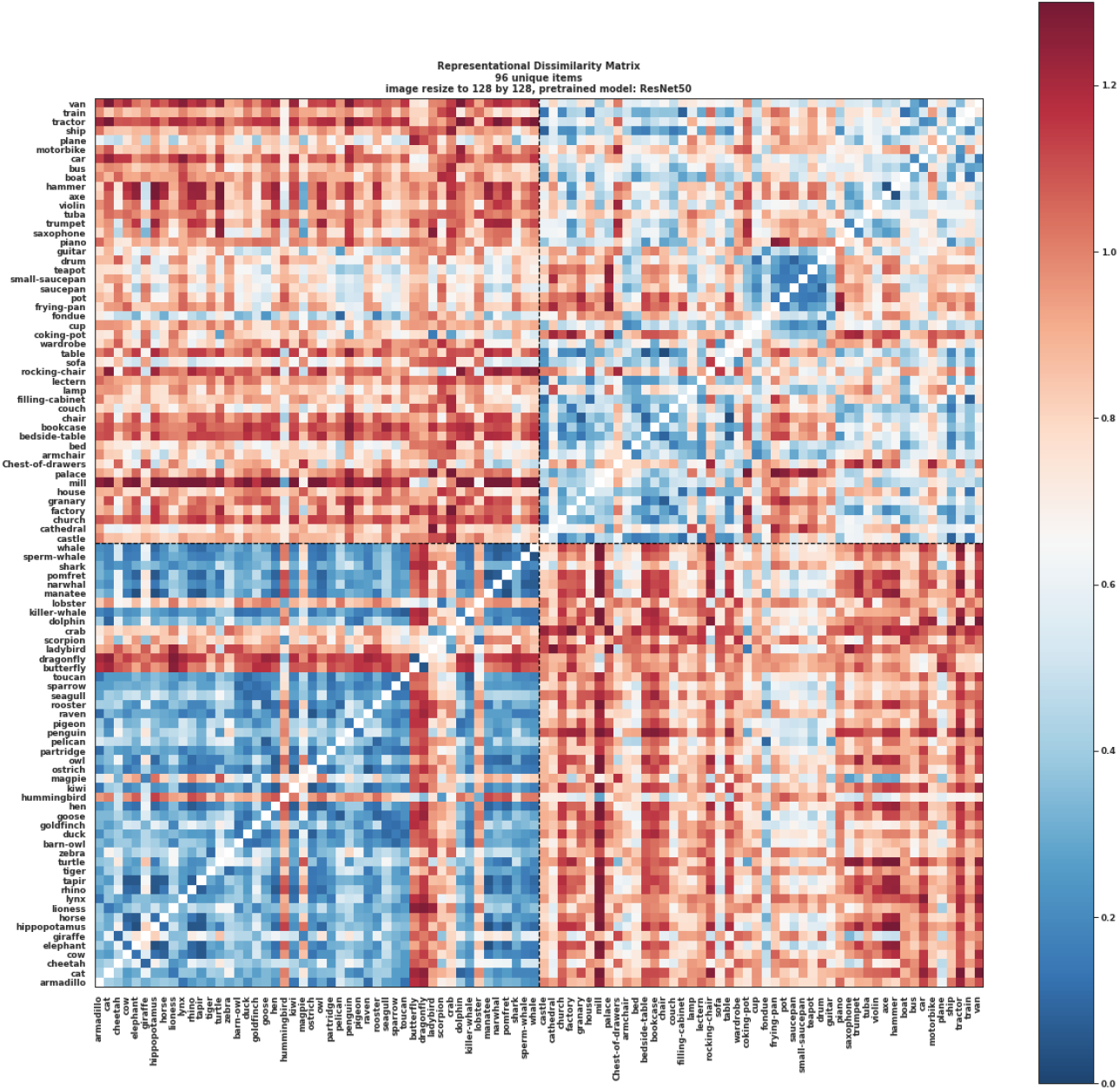
Representational dissimilarity matrix of the hidden representations of the Resnet50 model fined-tuned by Caltech101 dataset.

**Supplementary Figure 14:**
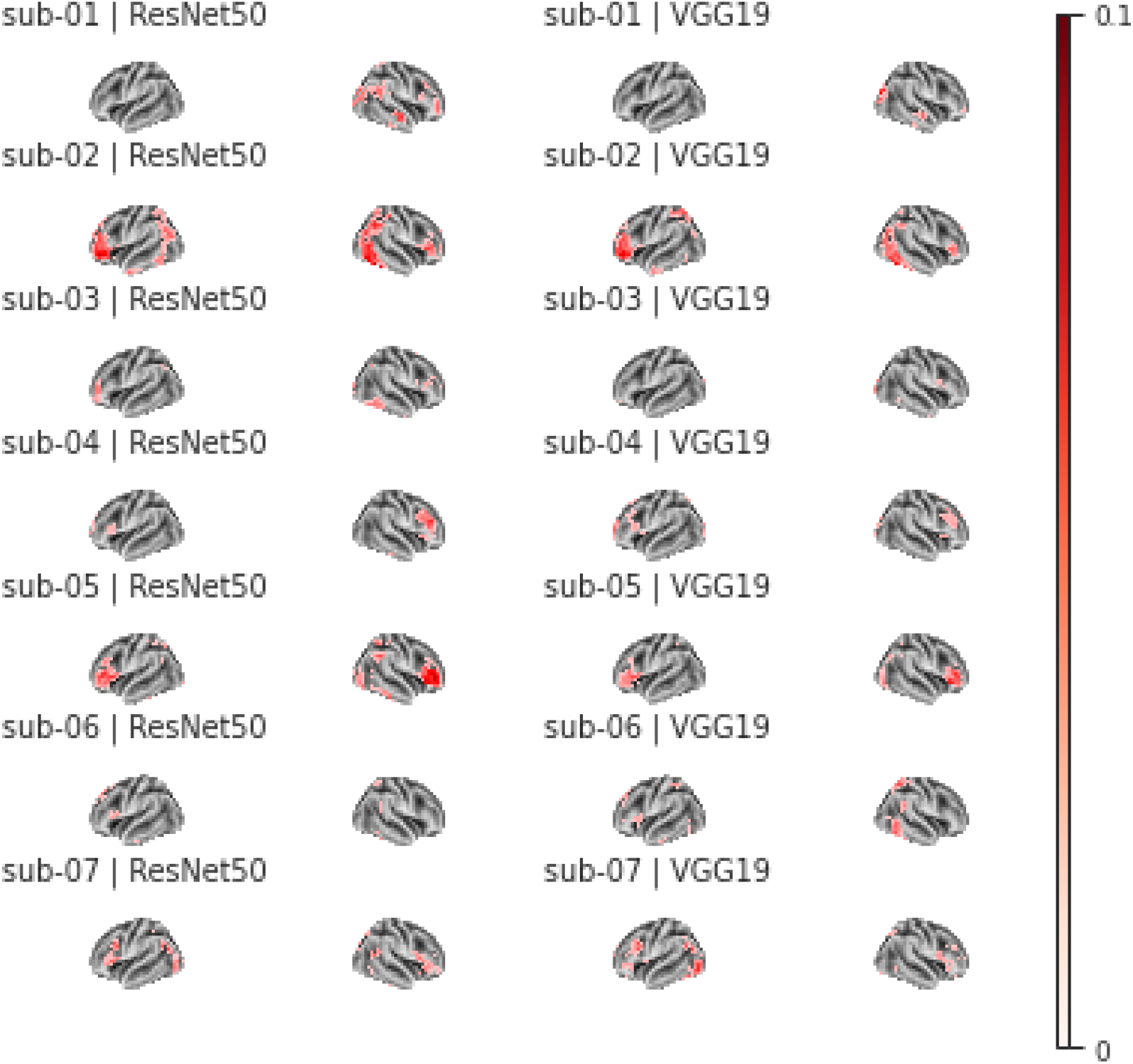
Searchlight representational similarity analysis in the unconscious trials illustrating the Spearman correlation between the model RDM and the fMRI-RDM.

**Supplementary Figure 15:**
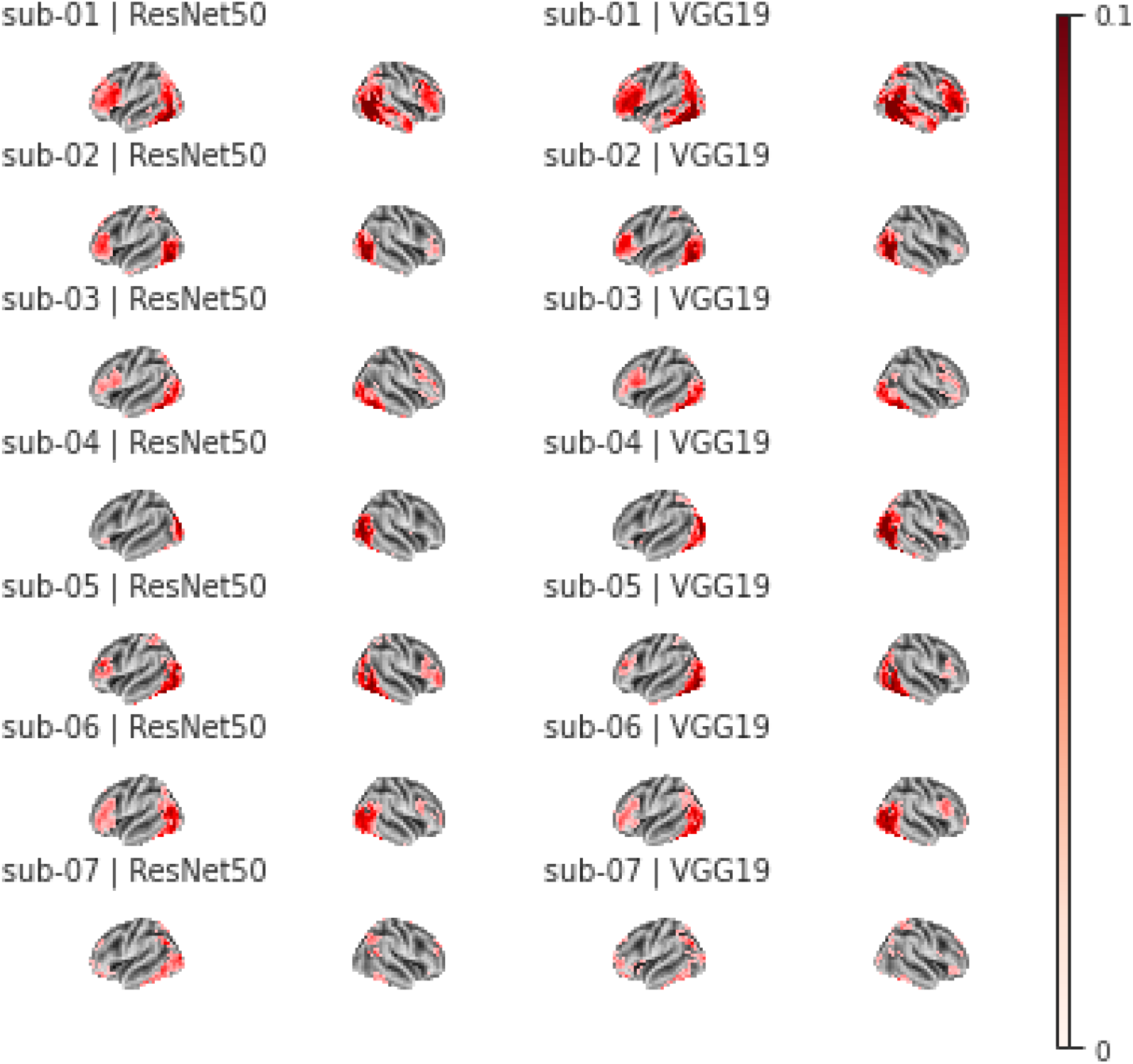
Searchlight representational similarity analysis in the conscious trials illustrating the Spearman correlation between the model RDM and the fMRI-RDM.

**Supplementary Figure 16:**
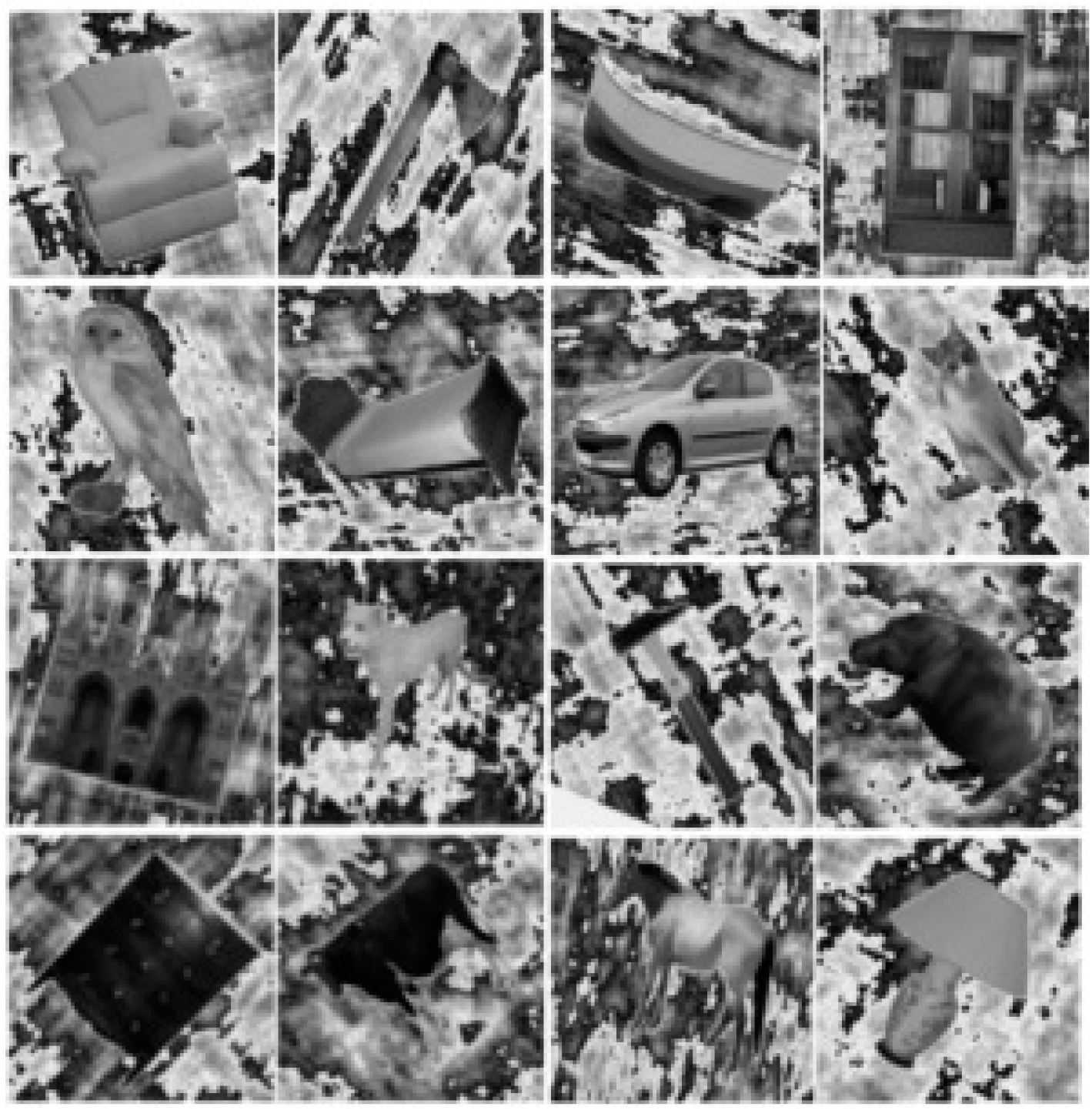
Examples of the images used in the fMRI experiment^5^.

**Supplementary Figure 17:**
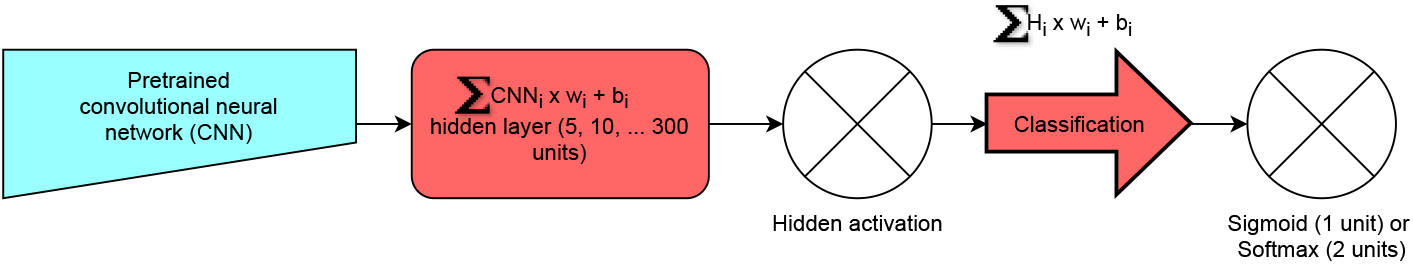
A simplified scheme of fine-tuning a pre-trained feedforward convolutional neural network model. The task is to classify the living vs. non-living category of the images (without noise) used in the fMRI experiment. The blue architecture was frozen and the weights were not updated, while weights of the red architectures were updated during training.

**Supplementary Figure 18:**
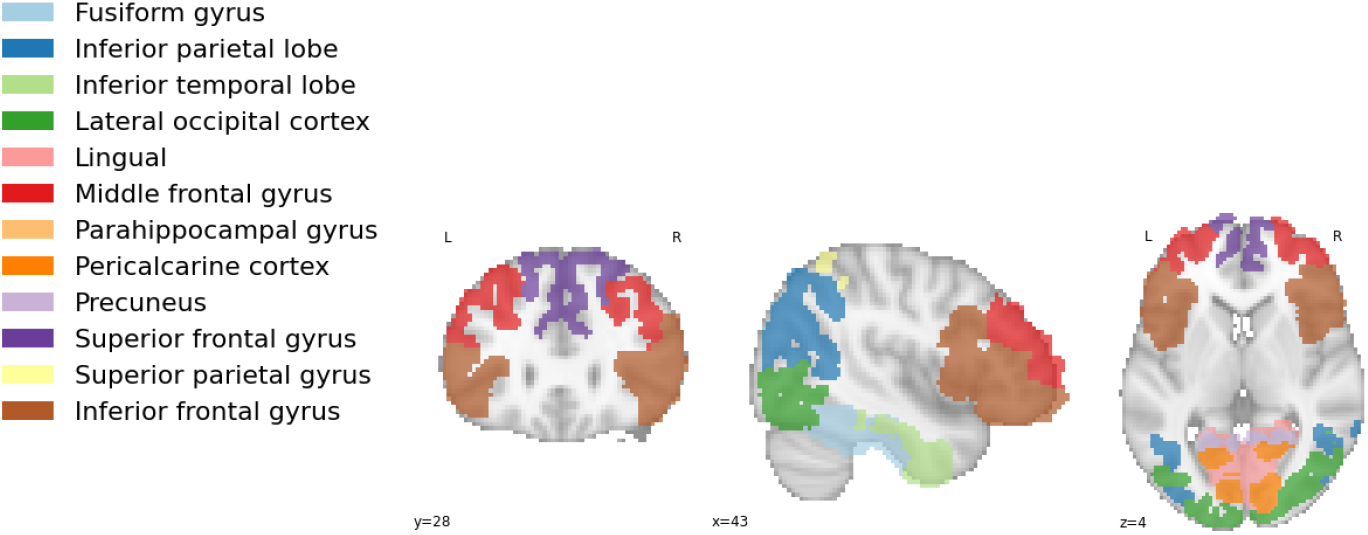
The figure shows the selected regions of interest. Twelve bilateral ROIs were extracted, comprising the lingual gyrus, pericalcarine cortex, lateral occipital, fusiform, parahippocampal cortex, precuneus, inferior temporal lobe, inferior and superior parietal lobe, superior frontal, middle frontal gyrus, and inferior frontal gyrus.

## Footnotes

a High definition figure: https://tinyurl.com/46yfyn92

b https://scikit-learn.org/stable/modules/generated/sklearn.model_selection.permutation_test_score.html

c https://tinyurl.com/up2txma

## Notes

### Competing Interest Statement

The authors have declared no competing interest.

### Summary of Updates

Included additional supplementary information with extended summary statistics.

